# A Whole-Plant Monocot from the Early Cretaceous

**DOI:** 10.1101/302075

**Authors:** Zhong-Jian Liu, Li-Jun Chen, Xin Wang

**Affiliations:** College of Landscape Architecture, Fujian Agriculture and Forestry University, Fuzhou 350002, China; Shenzhen Key Laboratory for Orchid Conservation and Utilization, National Orchid Conservation Center of China and Orchid Conservation & Research Center of Shenzhen, Shenzhen 518114, China; CAS Key Laboratory of Economic Stratigraphy and Palaeogeography, Nanjing Institute of Geology and Palaeontology, CAS, Nanjing 210008, China

**Keywords:** flower, fossil, Lower Cretaceous, China, angiosperms, herbaceous monocot

## Abstract

The Yixian Formation (the Lower Cretaceous) of China is world famous for its fossils of early angiosperms. Although these diverse angiosperms demonstrate an unexpectedly great diversity, few are preserved as whole plants (not mention of monocots), making our understanding of them incomplete. Here, we report a fossil angiosperm, *Sinoherba ningchenensis* gen. et sp. nov (Sinoherbaceae fam. nov.), from the Yixian Formation of China; this fossil has a physically connected underground stem with fibrous rootlets, a stem with branches and nodes, leaves with parallel-reticulate veins, and a panicle of female flowers with an ovary surrounded by perianth. Morphological and phylogenetic analyses revealed that *Sinoherba* is an herbaceous monocot taxon. This newly discovered fossil underscores the great diversity of angiosperms in the Lower Cretaceous Yixian Formation.

---

Although angiosperms constitute the most diversified plant group in the current world^*1*^, the origin, evolution and systematics of angiosperms are little understood or misunderstood. Early angiosperms including the famous *Archaefructus*^*1*-*4*^ have been reported from the Yixian Formation of northeastern China, but few of them are preserved as whole plants. Such fragmentary information hinders our full understanding of early angiosperms. To improve our understanding of early angiosperms, here, we report a new fossil plant, *Sinoherba ningchengensis* gen. et sp. nov. (Sinoherbaceae fam. nov.), from the Yixian Formation (the Early Cretaceous, 125 Ma). This fossil has a physically connected underground stem with nodes and fibrous rootlets; a stem with nodes, leaves and branches at the nodes; leaves with parallel-reticulate veins; a panicle arising from the axils of the leaves; and female flowers with an ovary surrounded by perianth. Morphological and phylogenetic analyses revealed that *Sinoherba* is an herbaceous monocot. Furthermore, *Sinoherba* demonstrates a novel character assemblage that helps elucidate the derivation of both gynoecia and growth habits of early angiosperms, underscores the great diversity of angiosperms in the Lower Cretaceous Yixian Formation, and suggests an earlier-than-assumed origin of angiosperms.

The Yixian Formation of China is well known for the Johel Biota as well as various fossil animals^*5*-*12*^ and plants^*13*-*17*^. Among the plants are Bryophyta, Lycopodiales, Equisetales, Filicales, Pteridospermae, Cycadales, Bennettitales, Ginkgoales, Czekanowskiales, Coniferales, and Angiospermae^*13*-*17*^. A general consensus exists with respect to the age of the Yixian Formation, namely, approximately 125 Ma (the Barremian-Aptian, Lower Cretaceous)^*19*^. The new fossil plant was collected from an outcrop of the Yixian Formation in Dashuangmiao, Ningcheng, Inner Mongolia, China (Fig. 1). The specimen was preserved as a compression/impression in thin-layered siltstone, and embedded coalified residue was present. The specimen is 26 cm long and 5 cm wide; it is preserved on a slightly yellowish grey siltstone slab approximately 25 cm × 38 cm (Fig. 2). The specimen was imaged using a Nikon D200 digital camera and a Nikon SMZ1500 stereomicroscope equipped with a Nikon DS-Fi1 digital camera. The distal portion of the plant was observed using a Leo 1530 VP scanning electron microscope (SEM) at Nanjing Institute of Geology and Palaeontology, CAS (NIGPAS). Sketches of the images were drawn using Photoshop 4.0 software.

**Figure 1.**
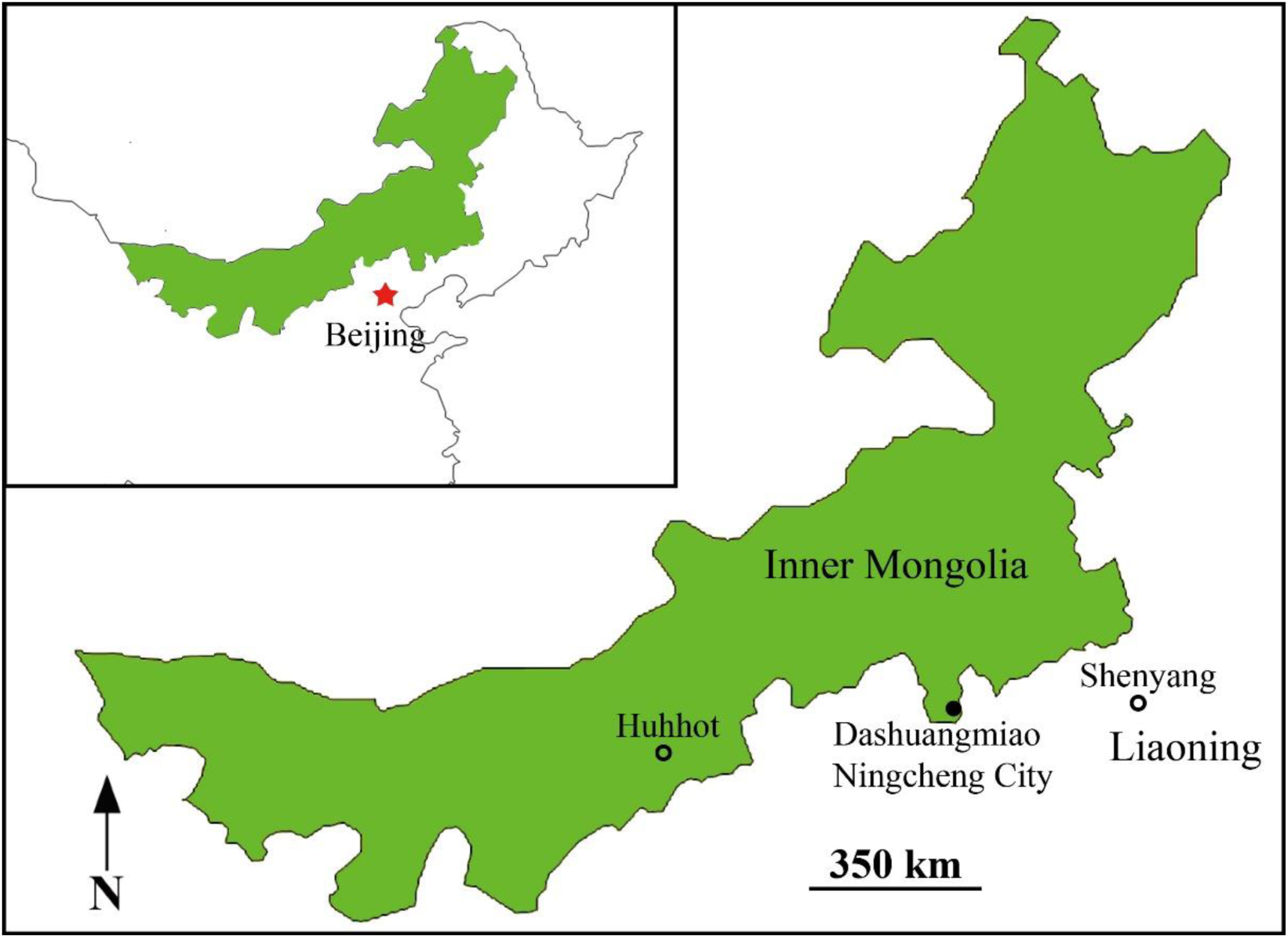
Fossil locations. The map shows the Dashuangmiao locality in Ningcheng of Inner Mongolia from where the fossil *Sinoherba ningchengensis* gen. et sp. nov. was collected. The inset map shows the location of Inner Mongolia in northeastern China (the asterisk indicates Beijing).

**Figure 2.**
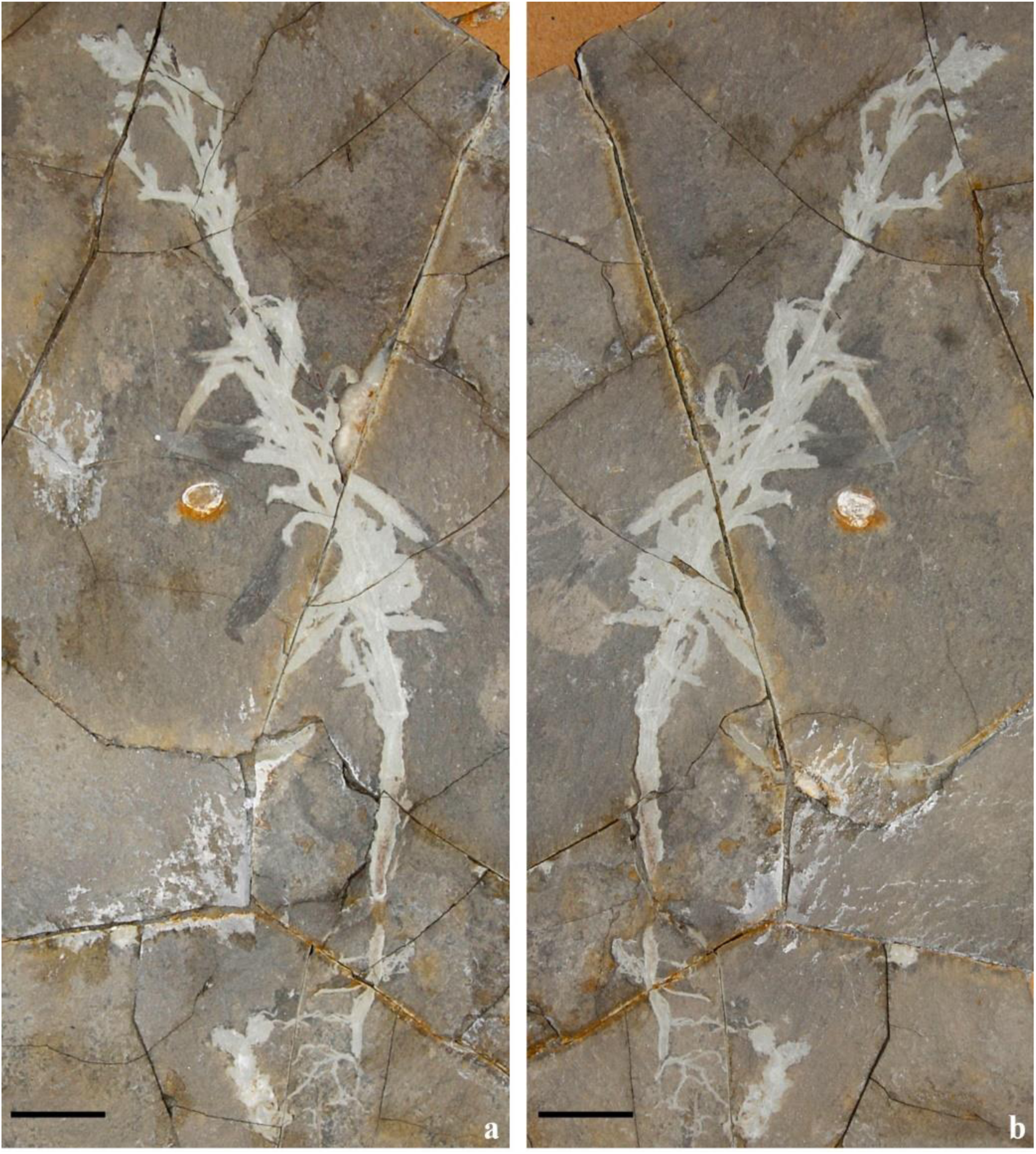
General view of the holotype of *Sinoherba* gen. nov. The whole plant includes the rootlets, stem, leaves and inflorescences. Bar = 2 cm.

**Angiospermae**

**Monocotyledoneae**

**Family: Sinoherbaceae** fam. nov.

**Genus: *Sinoherba*** gen. nov.

**Type species: *Sinoherba ningchengensis*** sp. nov.

**Family and generic diagnosis:** Herb, approximately 26 cm tall, with an underground stem. Nodes on the underground stem with lateral roots. Rootlets fibrous. Stem straight, with obvious nodes, with branches on the upper nodes. Leaves attached to the nodes in whorls, strap-like, with two orders of parallel veins and sparse meshes. Panicle arising from the axil of a leaf. Flowers dense on rachis, probably dioecious, pistil surrounded by 2–3 whorls of perianth. Ovary with a short distal style, partially separated in the distal half by a septum. Ovule basal, on the bottom of the ovary. Fruits ovate.

**Etymology:** *Sino*-Latin word for China, *-herba* from Latin word for herbaceous plant.

**Species:** *Sinoherba ningchengensis* sp. nov.

**Specific diagnosis:** the same as that of the family and genus.

**Description:** Whole plant including the roots, stem, leaves, and flowers, 258 mm long and 46 mm wide (Figs. 2–4). The underground stem is 3.4 mm in diameter and 39 mm in length, with several nodes (Fig. 3a). Fibrous roots are borne on the apex and nodes of underground stem, roots with 2–3 orders of fibrous rootlets (Figs. 3a, b). The stem is approximately 5 mm in diameter at the bottom, tapering to 1.4 mm distally, with evident and slightly swollen nodes (Figs. 2, 3c-e). The length of internodes varies from 7 to 27 mm (Figs. 2, 3c). Lateral branches/inflorescences and leaves are inserted on the nodes along the stem (Figs. 2, 3c-e, 4e). Younger lateral branches are in the axils of leaves (Figs. 2, 3e). The leaves are more concentrated on the basal half of the plant and are fully absent on two basal-most nodes of the stem (Figs. 2, 3c, f, g, 4e). A leafless node has leaf scars (Fig. 3d). The leaves are strap-shaped, up to 3.6 mm wide and 39 mm long, base contracted into a petiole (Fig. 3f) clasping the stem (Fig. 3g), apex acute to obtuse (Fig. 3h). The leaf veins are mostly parallel, with two orders of veins and elongated sparse meshes (Figs. 3f, i). The plant is mature, with two panicles (Figs. 4a, e). One of inflorescences arises from the apical node of the stem, the inflorescence has many young flowers, approximately 5 cm in length, with a bract at the base of the scape, mostly concentrated in the distal portion of the plant (Figs. 4a, b). The bract of inflorescence is oblong-lanceolate, apex obtuse (Fig. 4c). The flowers are terminal, alternately arranged along the peduncle, with a superior ovary, 0.2 mm wide and 0.2 mm long, surrounded by perianths (foliar parts) (Figs. 2, 4a, b). Another inflorescence, whose scape arises in the axil of a leaf on the stem, approximately 3.5 cm in length (Fig. 4a). The inflorescence is approximately 3 cm long, with several flowers/fruits (Fig. 4e). The flowers (fruits) are densely arranged along the peduncle and have 2–3 layers of perianth, 6 mm long and 5 mm wide, pedicel short, approximately 1 mm long (Figs. 4e, f). No male parts are recognized. The gynoecium includes a conical ovary and a distal style and is surrounded by perianth (Figs. 4f–h). The ovary is unilocular proximally but bilocular distally, with a wall approximately 67 μm thick and a basal ovule (Figs. 4d, g, h). The style is stout, cylindrical, up to 372 μm in length and 155 μm in diameter (Fig. 4h). The ovule is basal, approximately 670-1000 μm wide and 460–590 μm high (Fig. 4g, h, 6c-d).

**Figure 3.**
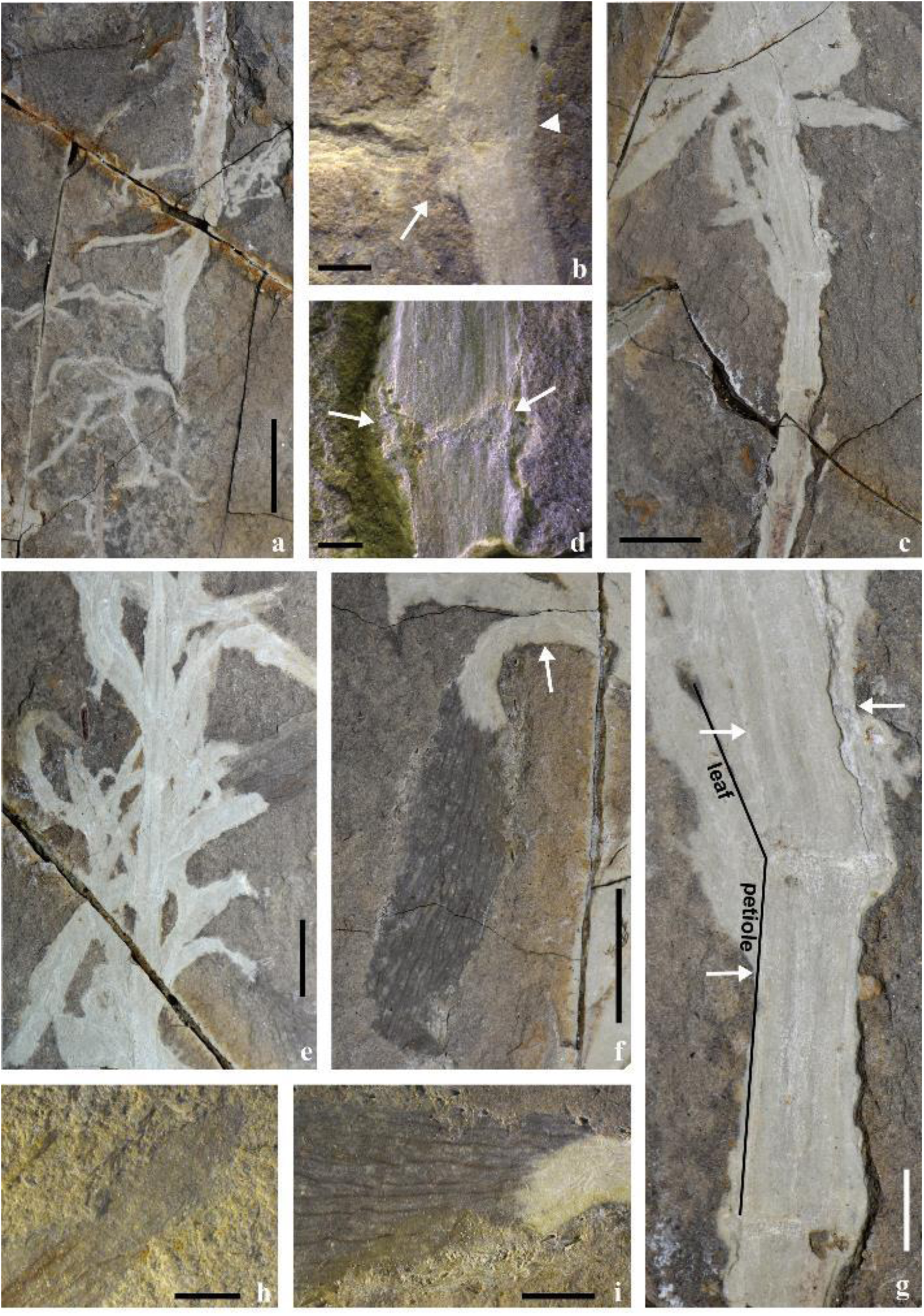
The vegetative organs of *Sinoherba* gen. nov. **a**. Fibrous roots are borne on the apex and nodes of an underground stem. Bar = 1 cm. **b**. A root (arrow) is borne on a node (arrowhead). Bar = 1 mm. **c**. A stem with swollen nodes and some nodes without leaves. Bar = 1 cm. **d**. A node with leaf scars (arrows). Bar = 1 mm. **e**. The stem together with a branch at the node. Bar = 1 cm. **f**. A leaf with parallel venation attached to a node. Note the leaf base that is contracted into the petiole (arrow). Bar = 1 cm. **g**. The petiole of a leaf clasping the stem (arrows). Bar = 3 mm. **h**. The apex of a leaf. Bar = 1 mm. **i**. Detailed view of the venation of the leaf shown in (f). Bar = 2 mm.

**Figure 4.**
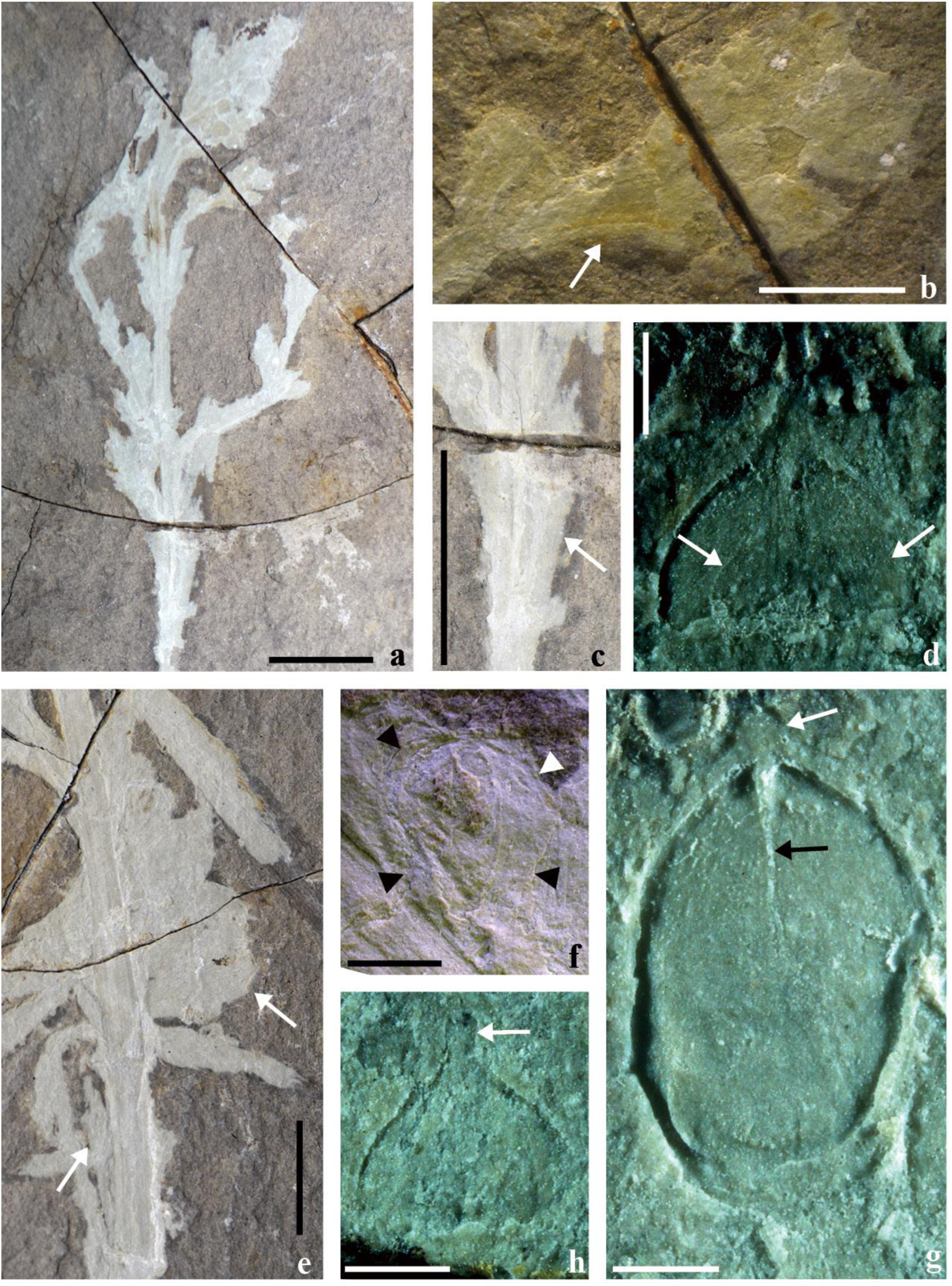
The reproductive organs of *Sinoherba* gen. nov. **a**. A panicle with many flowers. Bar = 1 cm. **b**. Flowers borne on a pedicel (arrow). Bar = 2 mm. **c**. A bract at the base of an inflorescence (arrow). Bar = 1 cm. **d**. An ovary with basal ovule (arrows). Bar = 0.5 mm. **e**. An inflorescence arisen from the axils of a leaf (arrows). Bar = 1 cm. **f**. A flower of the inflorescence from the axil of a leaf on the stem in Fig. 4e, showing the ovary surrounded by approximately three layers of perianths (arrowheads). Bar = 2 mm. **g**. An ovary split through its centre, showing its style (white arrow) and half septum (black arrow). Bar = 0.5 mm. **h**. Detailed view of the distal portion of an ovary, showing the distal style (arrow) on the ovary. Bar = 0.5 mm.

**Etymology:** *ningcheng*-, for Ningcheng City, Inner Mongolia, the fossil locality.

**Holotype:** LJNG0002 a & b (Fig. 2).

**Type locality:** Liujianangou, Dashuangmiao Town, Ningcheng, Inner Mongolia, China (41°30.3792′N, n8°51.2465′E, Fig. 1).

**Stratigraphic horizon:** the Yixian Formation, equivalent to the Barremian-Aptian, Lower Cretaceous (125 Ma).

**Depository:** The Orchid Conservation & Research Center of Shenzhen, China (NOCC).

The small size and maturity (suggested by the presence of flowers) of *Sinoherba* indicate that it is an herbaceous plant. To the best of our knowledge, herbaceous growth habit among living seed plants is restricted to angiosperms. Therefore, the herbaceous growth habit of *Sinoherba per se* suggests that this plant is likely an angiosperm^*18*^, a conclusion supported by other features of *Sinoherba.*

*Sinoherba* has a panicle subtended by an inflorescence bract, and the flower is composed of a pedicel, perianths and a gynoecium. Importantly, via their enclosed ovule before pollination, angiosperms are easily distinguished from gymnosperms; enclosed ovules before pollination have never been reported in gymnosperms^*19*,*20*^. As seen in Figs. 4d, f–h and 6c-d, the basal ovule is fully enclosed by an ovary, the latter of which is completely sealed at the tip by a short style and is partially bilocular distally (Figs. 4g, h). These features are characteristic of angiosperms. As such, the angiospermous affinity of *Sinoherba* suggested above is confirmed.

The leaves of *Sinoherba* are arranged at its nodes, and the leaf petioles clasp the stem (Figs. 2, 3c–i); the leaves are not spirally arranged as might be expected according to angiosperm evolution theories^*21*^. Spiral phyllotaxy was formerly assumed to be ancestral in angiosperms^*21*^, and unfortunately, this belief is still pervasive among botanists. However, the phyllotaxy of *Sinoherba* apparently contradicts this belief. *Sinoherba* presents leaf characteristics that suggest a monocot affinity. The leaves of *Sinoherba* are oblong-lanceolate and with reticulate veins. The veins are of two orders: the first-order parallel veins converge at the apex, and the second-order veins join the first-order ones (Figs. 3f, h, i). The venation of *Sinoherba* is approximately parallel, a venation rarely seen in eudicots. The underground stem with fibrous rootlets (Fig. 3a) excludes a possible Eudicot affinity for *Sinoherba*, as this type of stem morphology is frequently seen in monocots but rarely seen in eudicots^*22*^. Such a character assemblage and herbaceous habit suggest that *Sinoherba* is a monocot.

*Sinoherba* is probably an helobious plant. An herbaceous nature and roots borne on the basal nodes of underground stem suggest an helobious habitat. The absence of leaves and the presence of leaf scars on the basal nodes of the stem in *Sinoherba* (Figs. 3c, d) suggest that the leaves must have fallen off before fossilization. *Sinoherba* has two types of inflorescences: one is a developing panicle, and the other appears to be an infructescence that probably formed during a different flowering season. In contrast to the presence of leaves on the nodes discussed above, this phenomenon suggests that *Sinoherba* is likely a perennial and deciduous herb that can adapt to environment changes (suggested by leaf falling). Similar character assemblage has been reported for various monocot families, e.g., the Helobiae^*23*^.

To decipher the origin, evolution and systematics of angiosperms correctly and confidently, all hypotheses must be based on fossil evidence. Various fossils of early angiosperms from the Yixian Formation, including those of *Archaefructus*^*1*-*4*^ and *Sinocarpus*^*24*,*25*^, have shed important light on these issues, but few of those fossils are of whole plants. The fossil of *Sinoherba* is unique because it is an herbaceous whole plant, including the roots, stem, leaves, and inflorescences. Phylogenetic analyses of the morphological characters as well as the combination of characters and molecular data of both extant plants and *Sinoherba* demonstrate that *Sinoherba* is a monocot taxon^*27*^ (Figs. 5, S1, S2).

When all the morphological characters of *Sinoherba* are evaluated in a phylogenetic context, *Sinoherba* nests in the angiosperm clade. The angiosperm clade is divided into three subclades: *Archaefructus*, monocots and eudicots. *Archaefructus* is a basal clade and is the sister of the monocots and eudicots. The Amborellaceae is the most basal subclade among the eudicots, while the Nymphaeaceae is the most basal subclade among monocots. *Sinoherba* is a monocot taxon and is the sister of the Najadaceae, an aquatic plant family (Fig. S1). Phylogenetic relationships were reconstructed using a combination of morphological characters and plastid DNA regions. The phylogenetic relationships revealed in this analysis (Fig. 5, Fig. S2) are similar to those revealed via morphologic characters alone. All analyses consistently placed *Sinoherba* among the monocots and indicated that it represents a sister of the extant Araceae and Helobiae families (Najadaceae, Alismataceae and Hydrocharitaceae). We hope future studies can test whether *Sinoherba* stands for the basalmost monocot. The characters unique to the Sinoherbaceae are illustrated in reconstruction of *S. ningchengensis* (Fig. 6).

**Figure 5.**
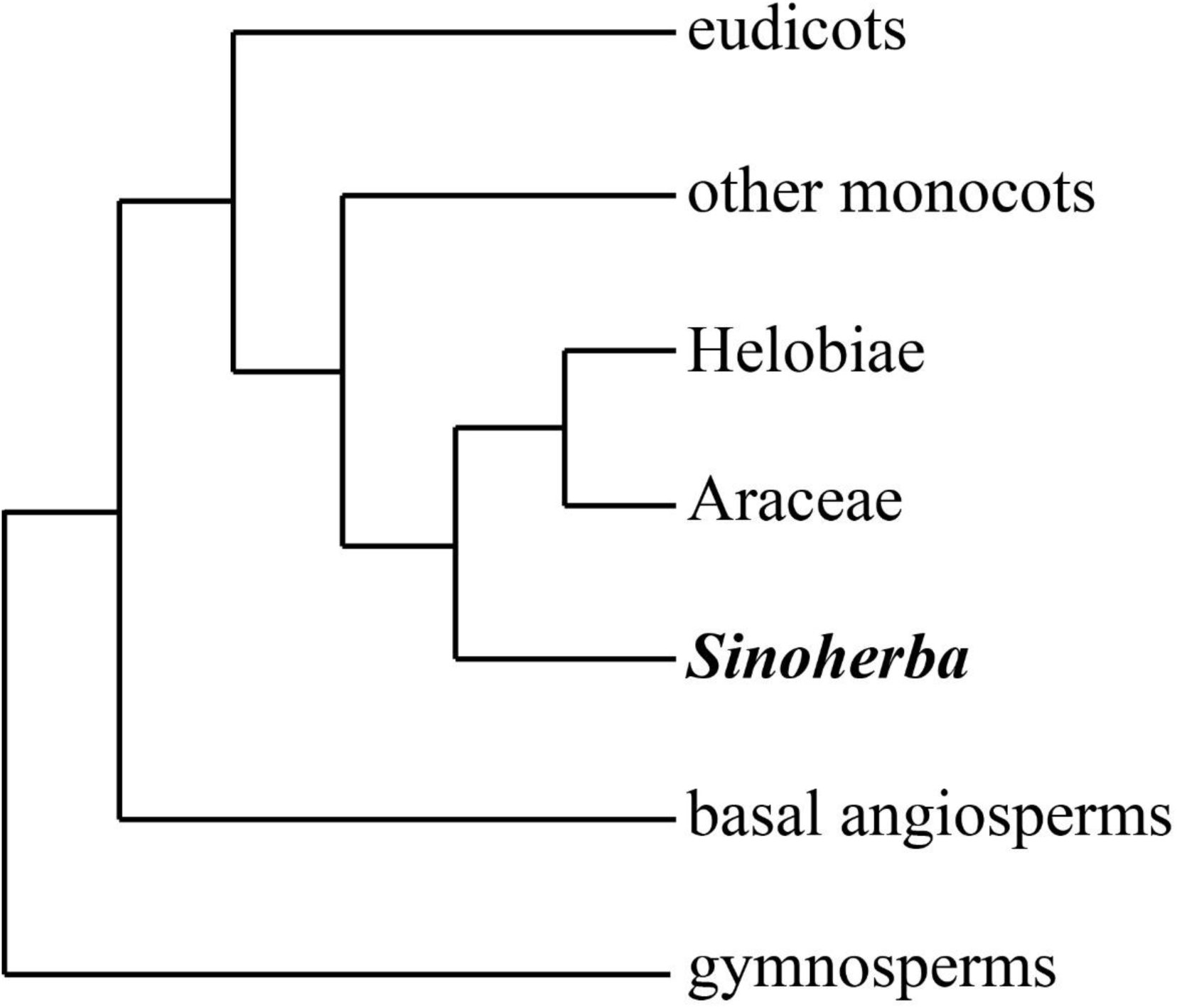
A phylogenetic tree (simplified) was reconstructed using a combination of morphological characters and plastid DNA regions (see details in Supplementary Fig. 2).

**Figure 6.**
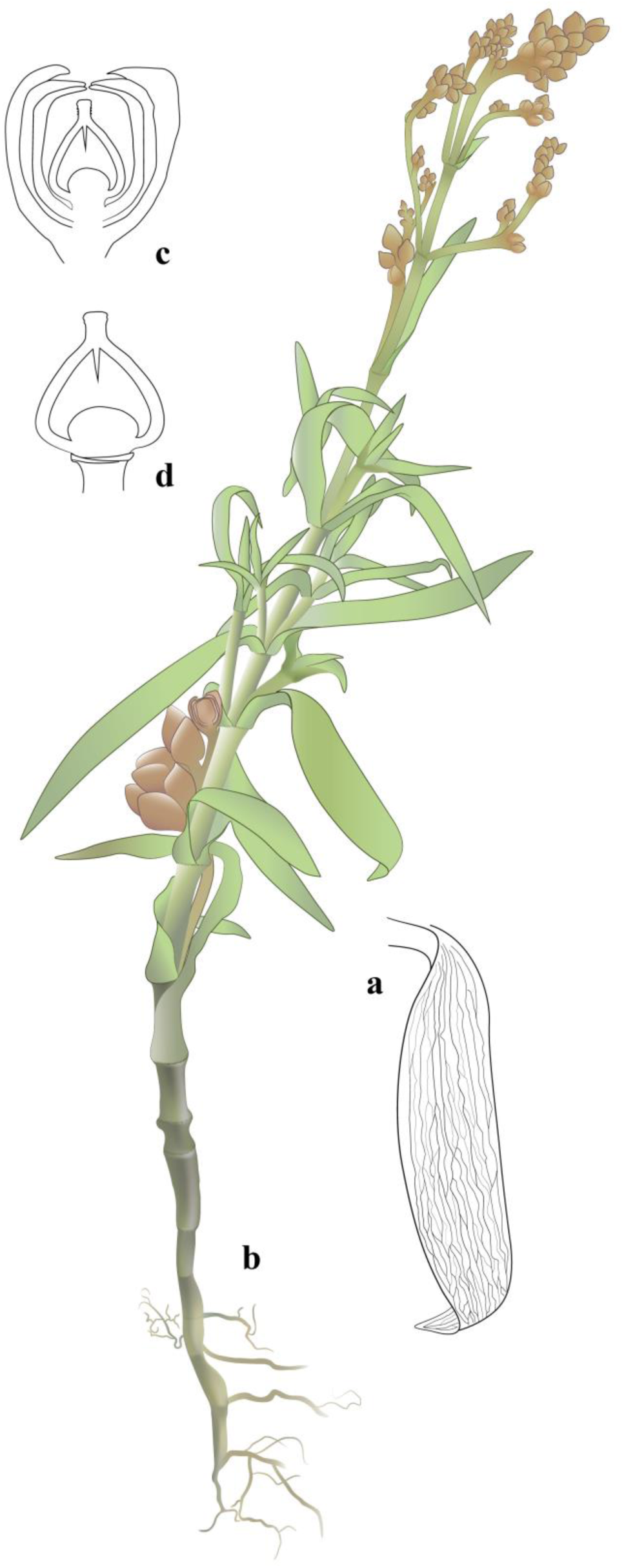
Reconstruction of *Sinoherba ningchengensis* gen et sp. nov. The plant is herbaceous and is composed of an underground stem with fibrous rootlets borne on the nodes, a stem with leaves (a) and branches in axils of bracts borne on the nodes, and inflorescences that both protrude from the axils of the leaves and are borne on the terminal stem (b). The pistil is surrounded by perianth (c) and the ovule on the bottom of the ovary (d). The colour of the flower was artificially created.

The occurrence of basal ovule in *S. ningchengensis* is unexpected for classical angiosperm evolution. However, similar way of gynoecium forming is commonly seen in angiosperm and is recently termed as mixomery^*26*^. Apparently, this new findings in extant as well as fossil angiosperms deserve attention in future study of angiosperm evolution.

The discovery of *S. ningchengensis* and the phylogenetic and morphological analyses refute the Early Cretaceous origin of monocots. Instead, our results favour a more ancient origin of angiosperms.

## Supplementary Information

**Table S1.**
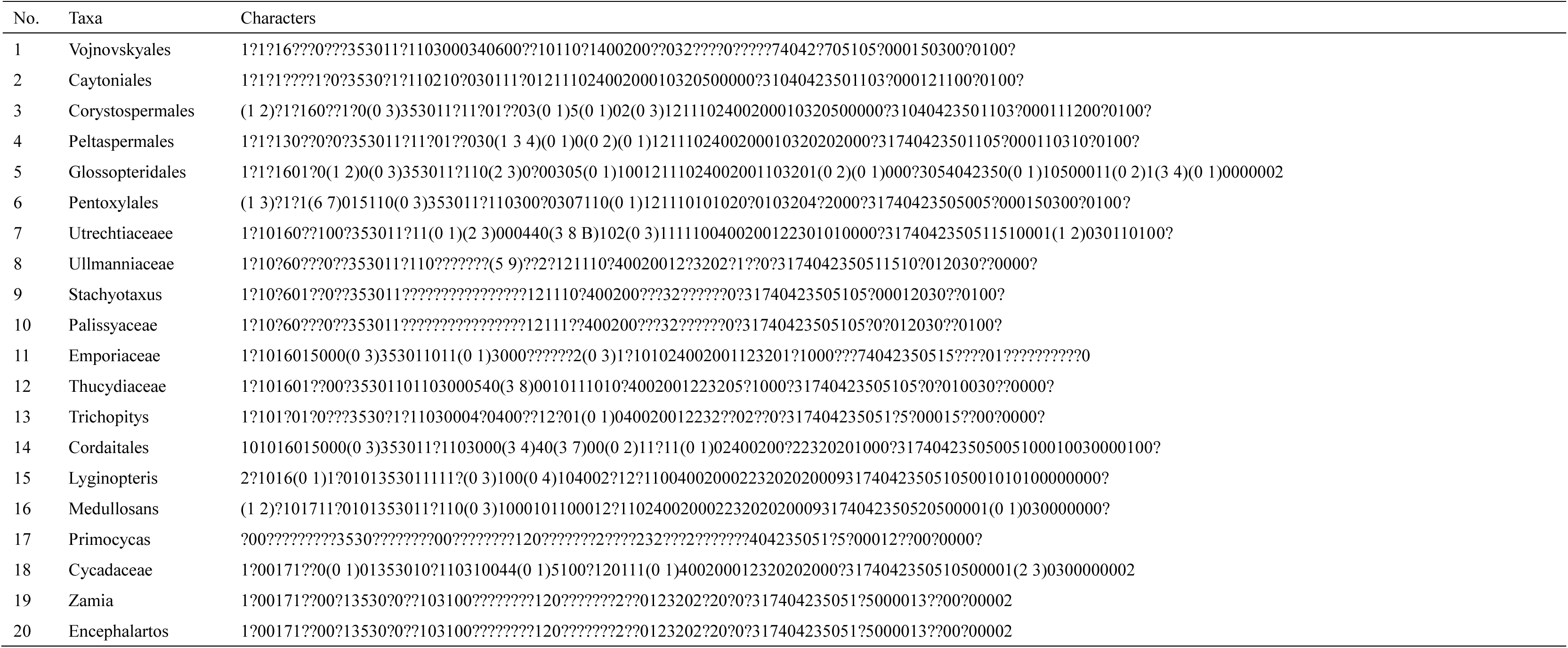
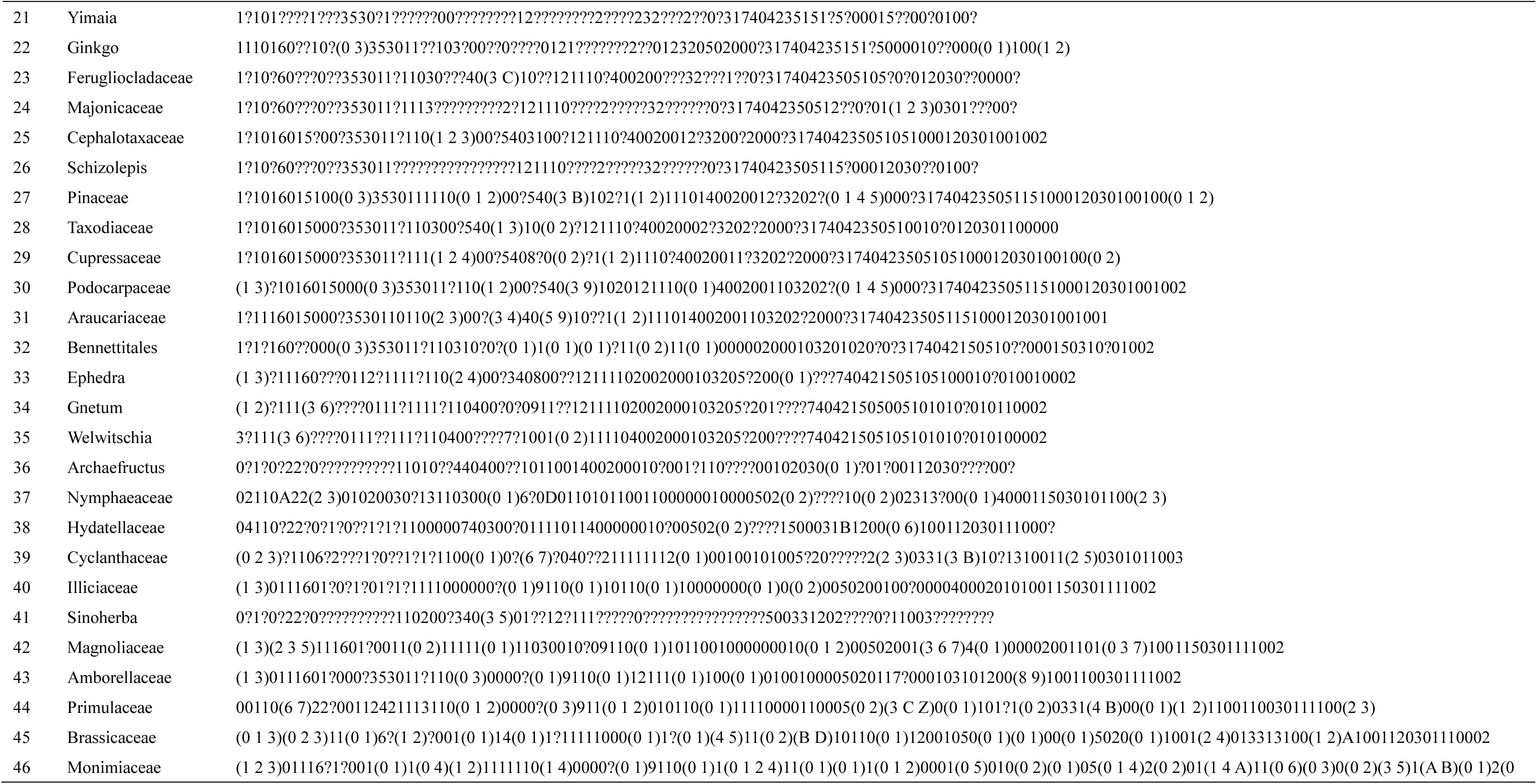
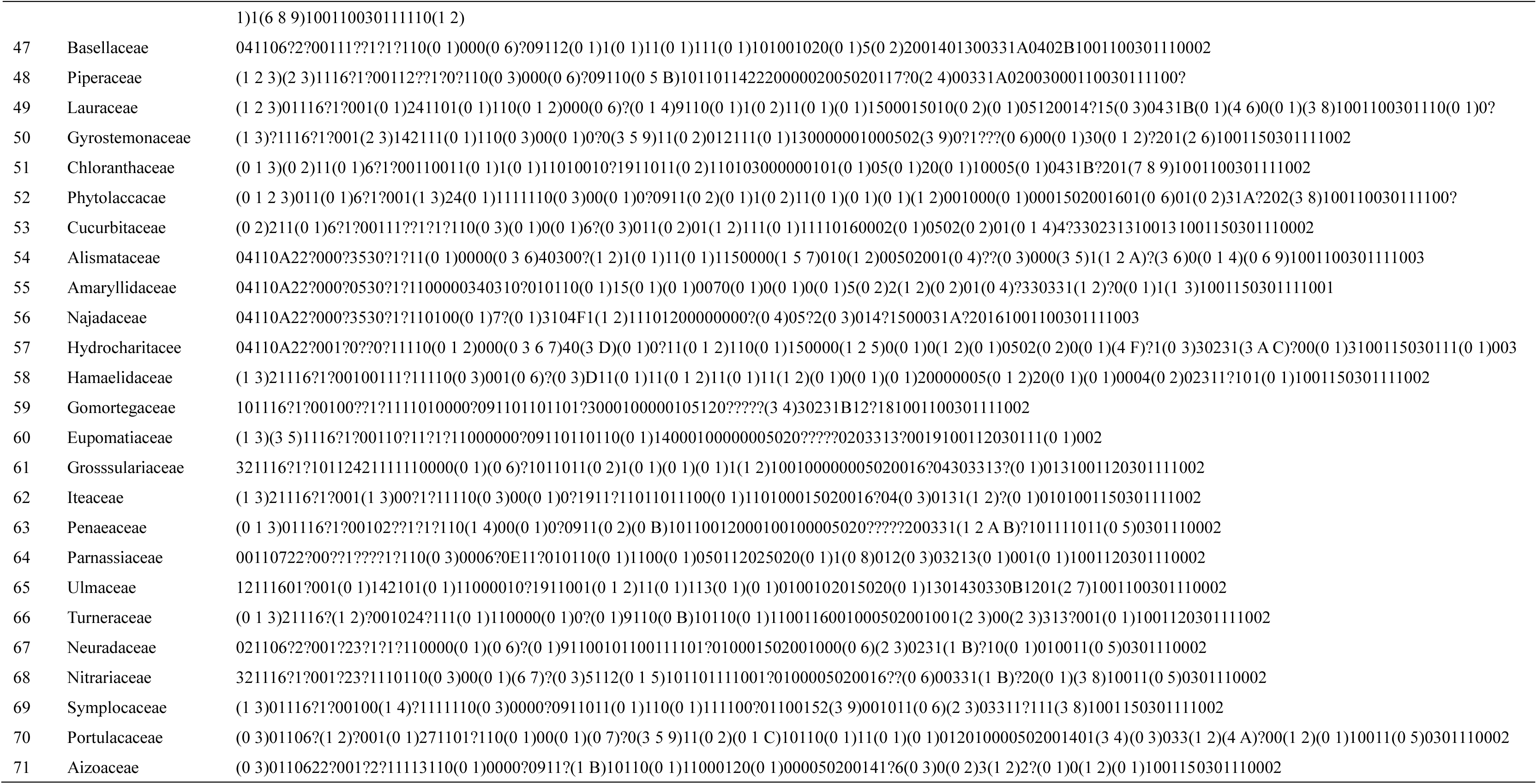
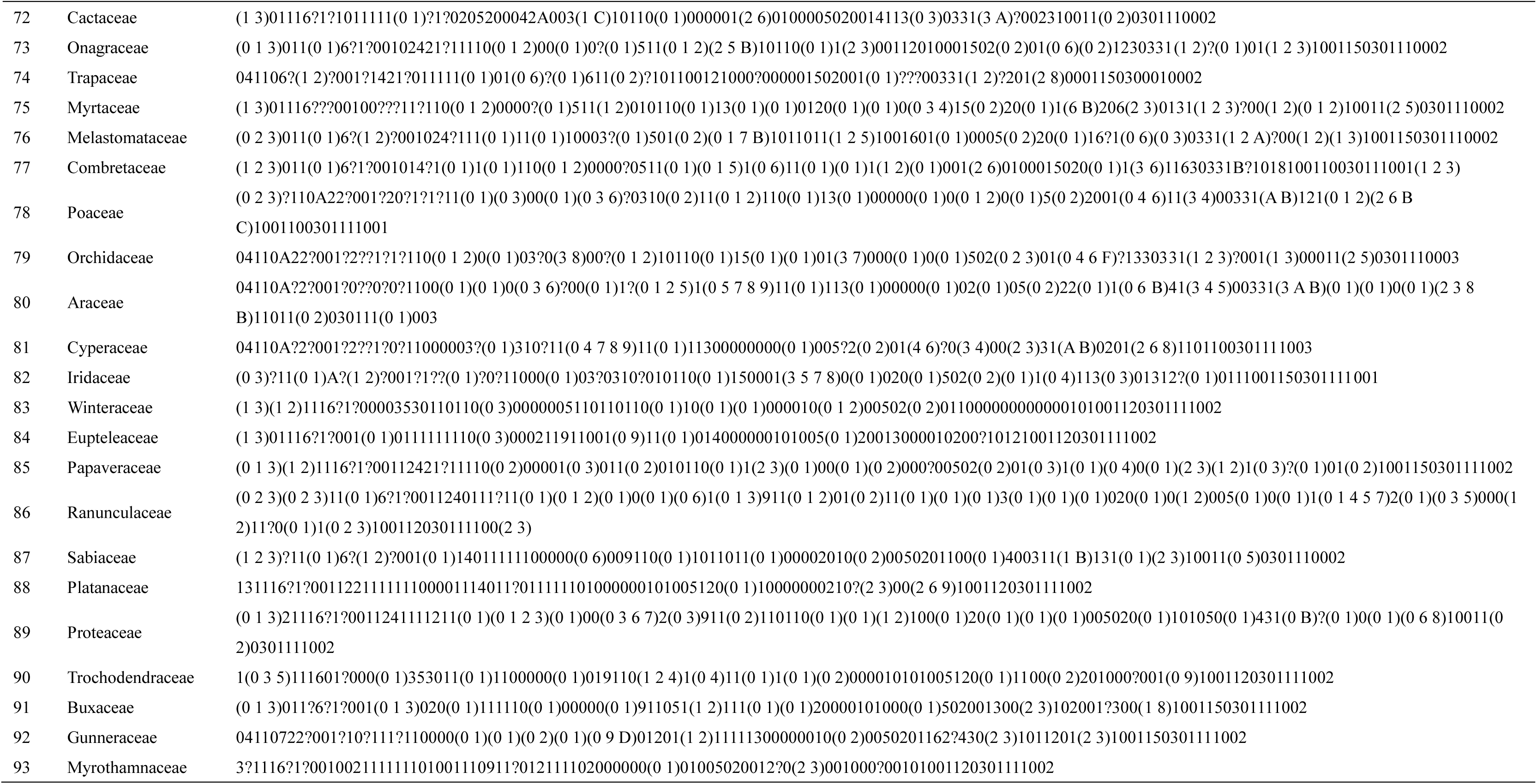
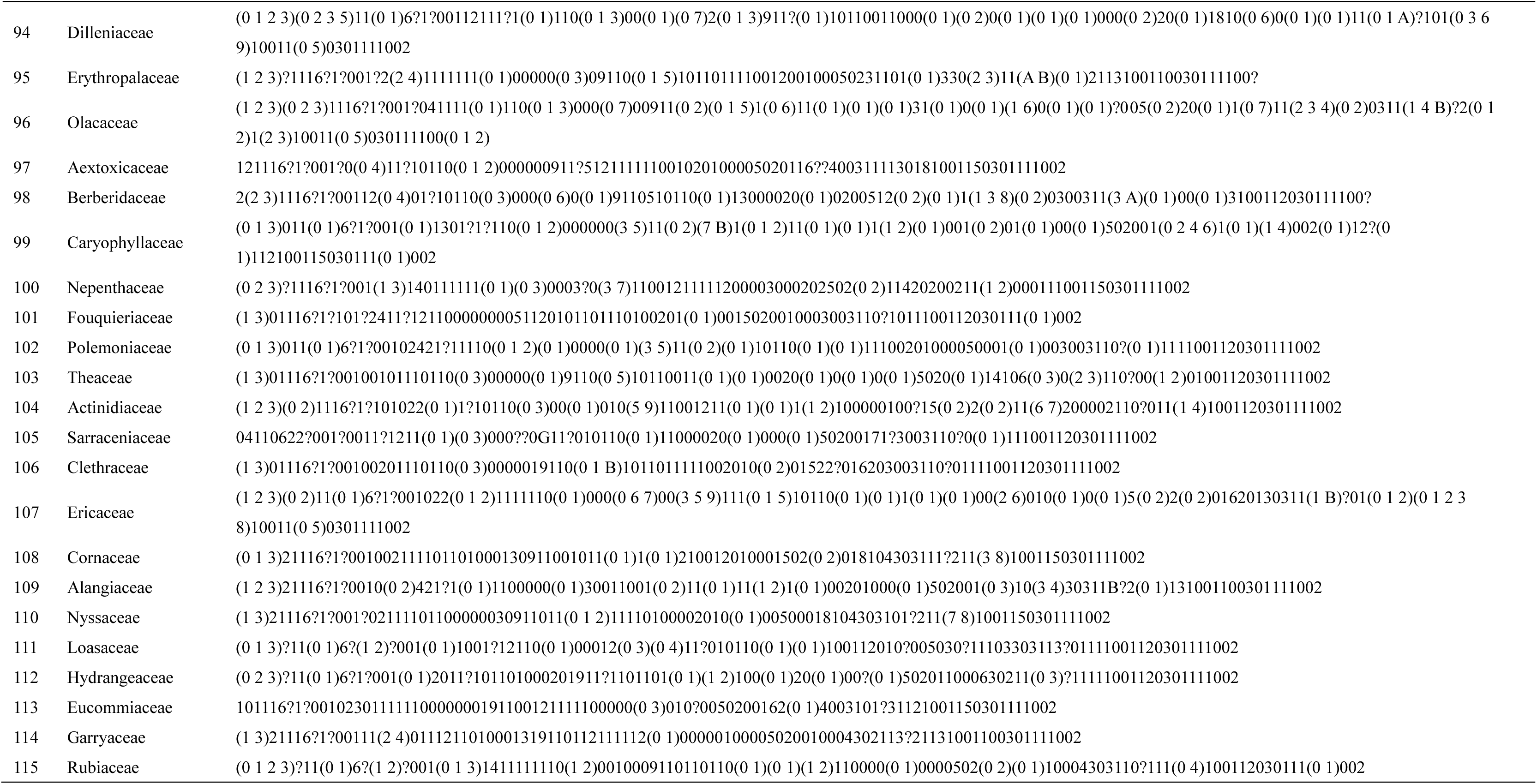
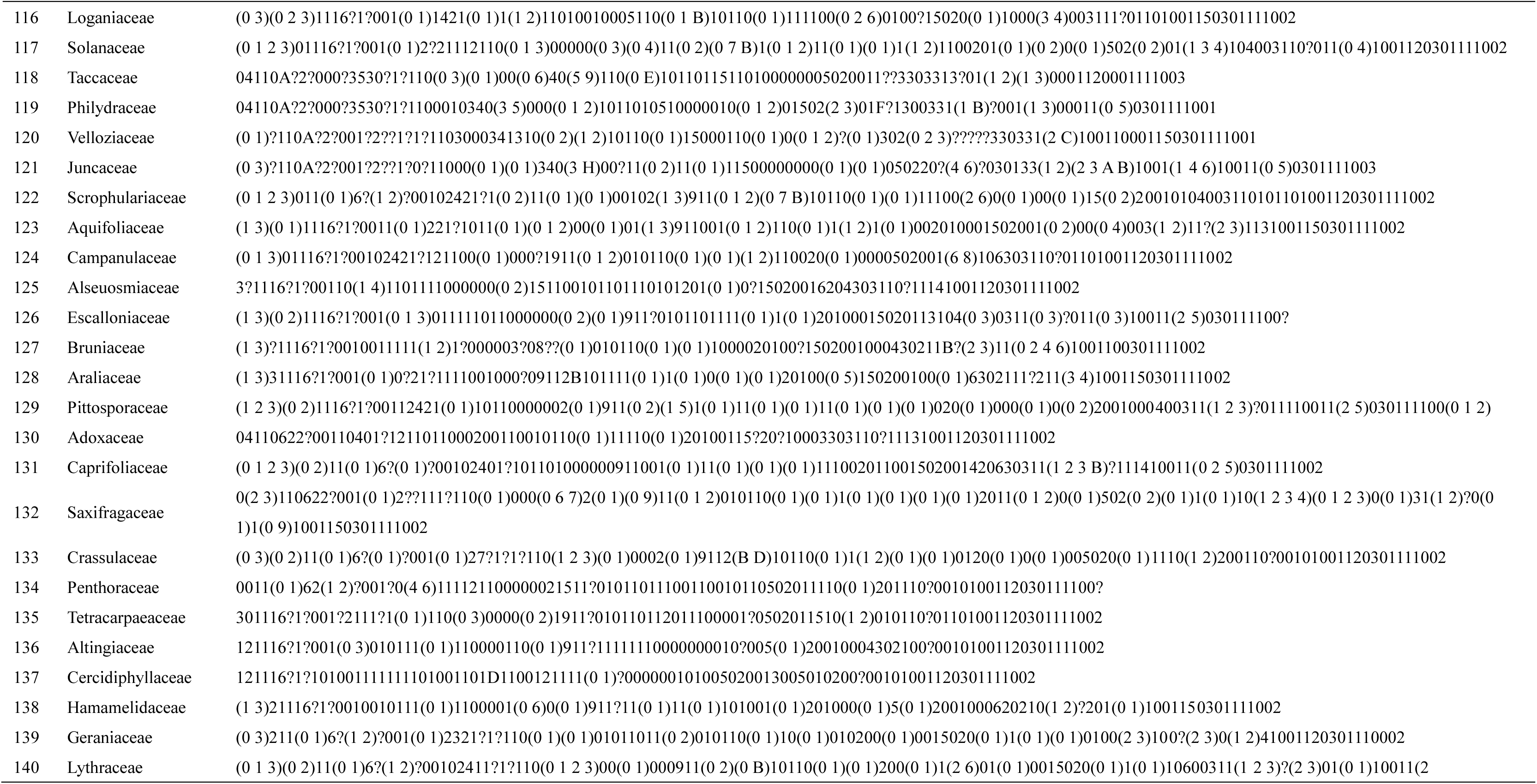
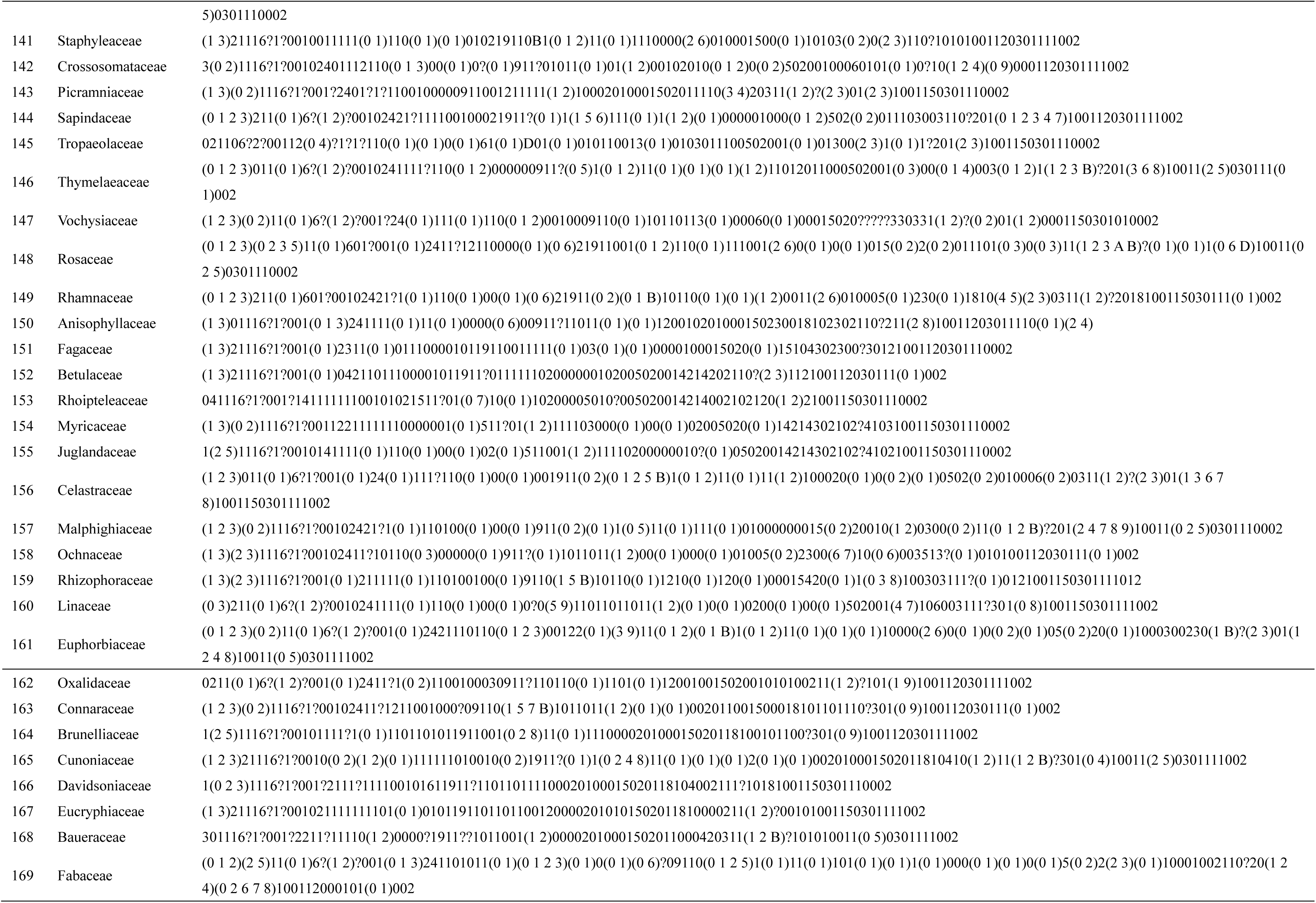
Taxa and value information.

**Table S2.**
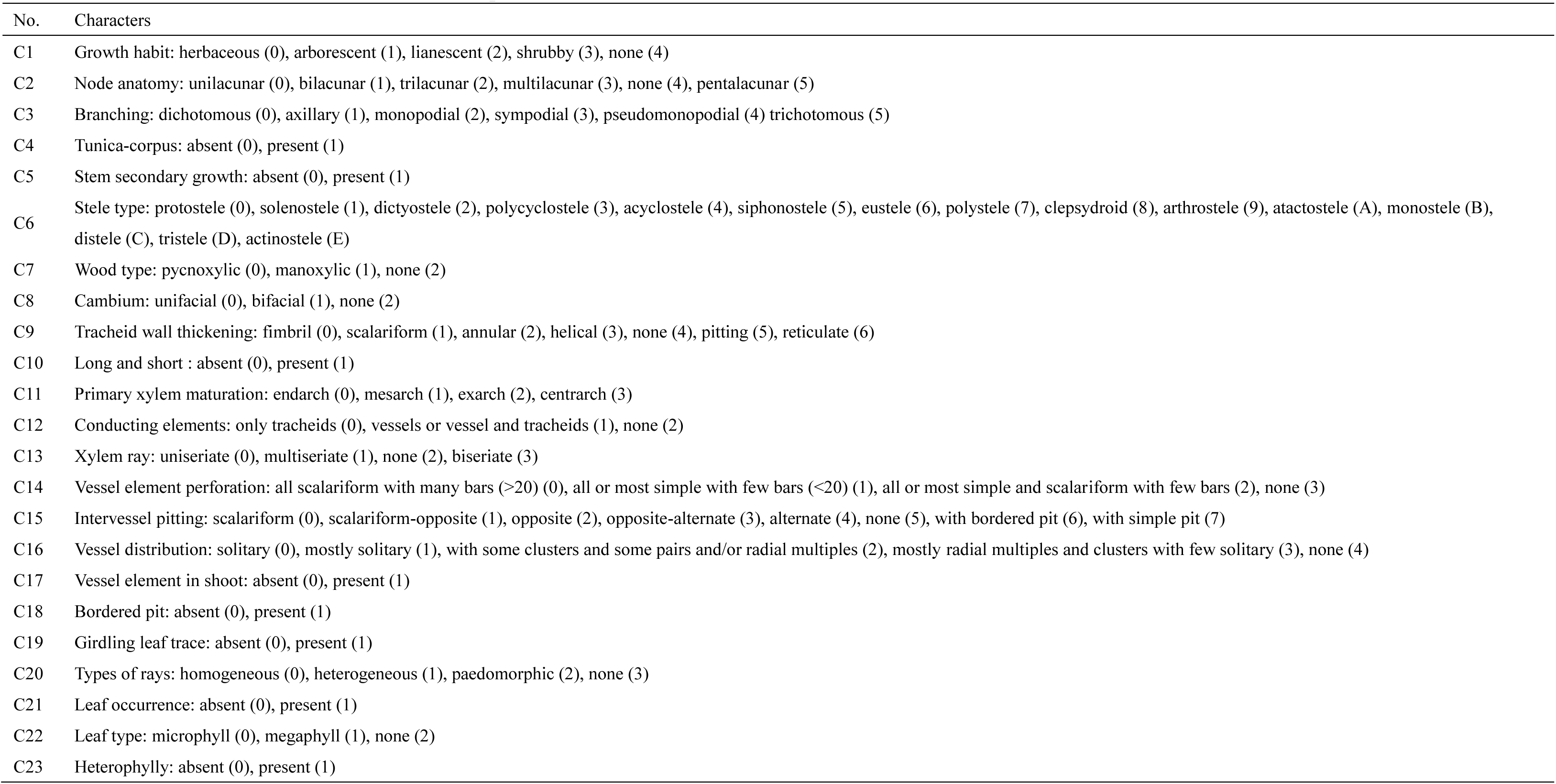
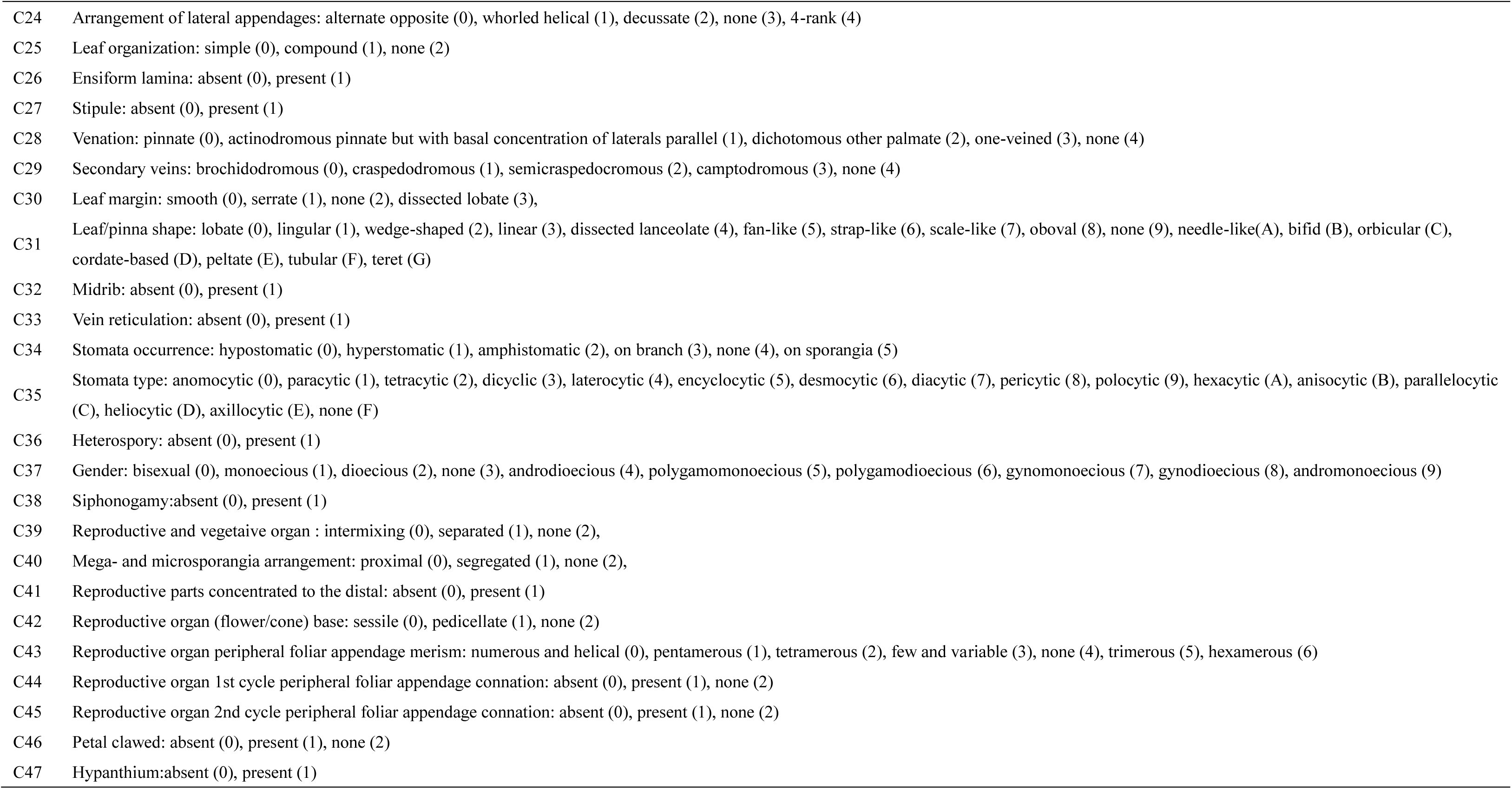
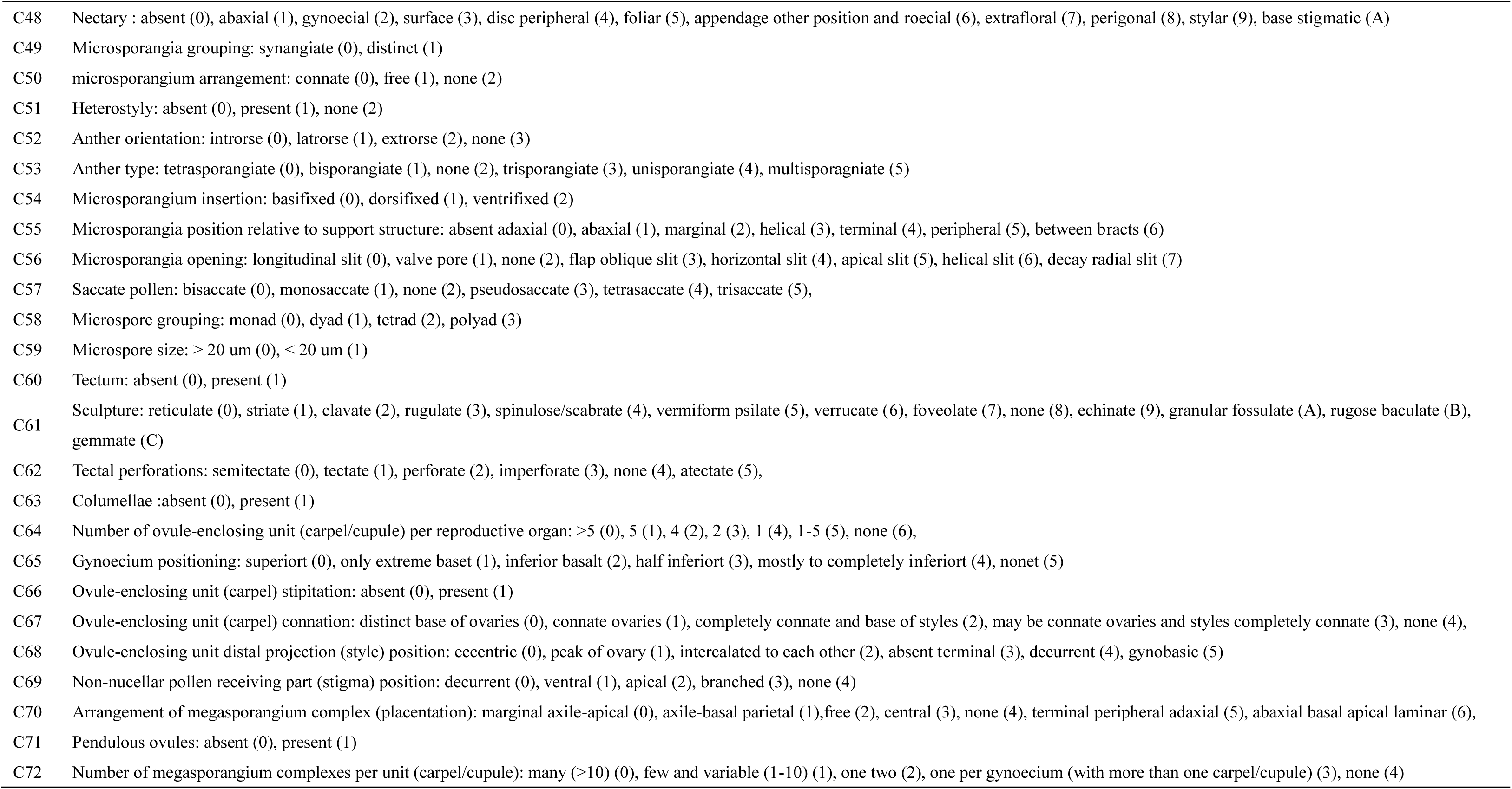
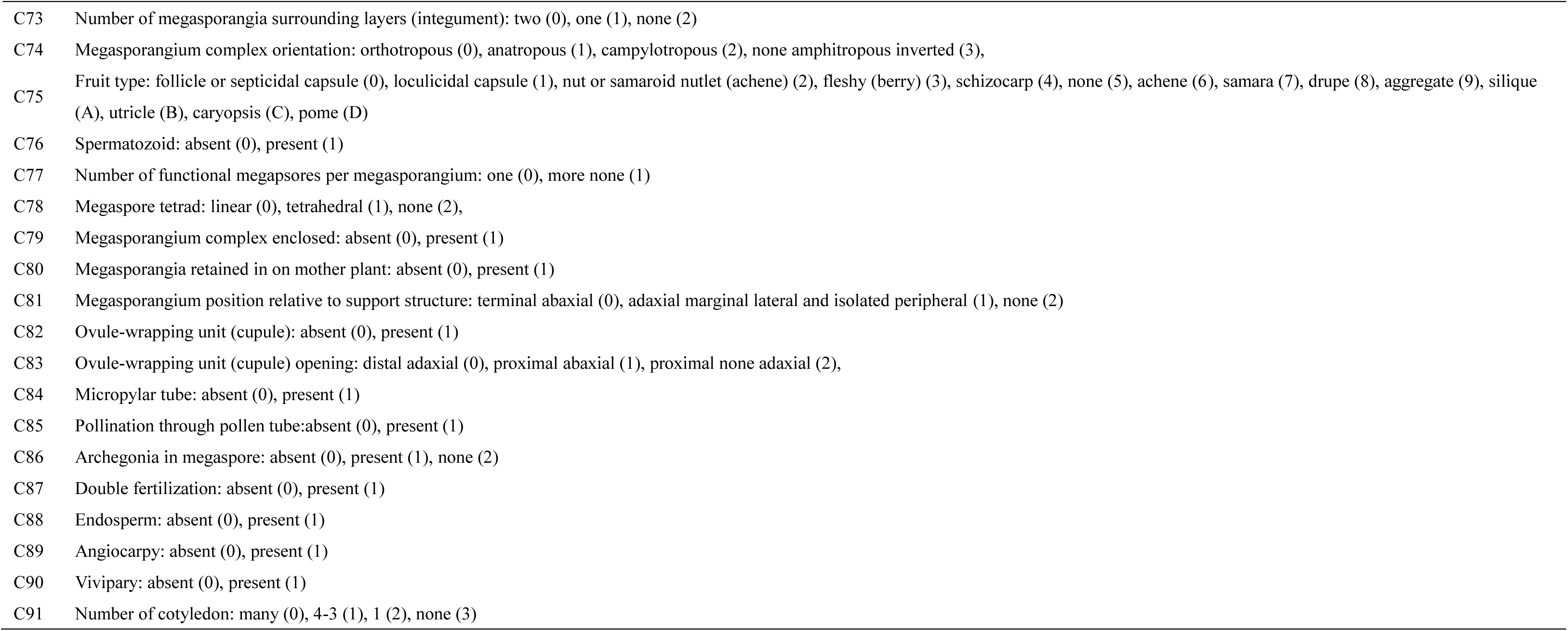
Characters (The numbers in the parentheses are the values of each characters).

| No. | Characters |
| --- | --- |
| C1 | Growth habit: herbaceous (0), arborescent (1), lianescent (2), shrubby (3), none (4) |
| C2 | Node anatomy: unilacunar (0), bilacunar (1), trilacunar (2), multilacunar (3), none (4), pentalacunar (5) |
| C3 | Branching: dichotomous (0), axillary (1), monopodial (2), sympodial (3), pseudomonopodial (4) trichotomous (5) |
| C4 | Tunica-corpus: absent (0), present (1) |
| C5 | Stem secondary growth: absent (0), present (1) |
| C6 | Stele type: protostele (0), solenostele (1), dictyostele (2), polycyclostele (3), acyclostele (4), siphonostele (5), eustele (6), polystele (7), clepsydroid (8), arthrostele (9), atactostele (A), monostele (B), distele (C), tristele (D), actinostele (E) |
| C7 | Wood type: pycnoxylic (0), manoxylic (1), none (2) |
| C8 | Cambium: unifacial (0), bifacial (1), none (2) |
| C9 | Tracheid wall thickening: fimbrial (0), scalariform (1), annular (2), helical (3), none (4), pitting (5), reticulate (6) |
| C10 | Long and short : absent (0), present (1) |
| C11 | Primary xylem maturation: endarch (0), mesarch (1), exarch (2), centrarch (3) |
| C12 | Conducting elements: only tracheids (0), vessels or vessel and tracheids (1), none (2) |
| C13 | Xylem ray: uniseriate (0), multiseriate (1), none (2), biseriate (3) |
| C14 | Vessel element perforation: all scalariform with many bars (>20) (0), all or most simple with few bars (<20) (1), all or most simple and scalariform with few bars (2), none (3) |
| C15 | Intervessel pitting: scalariform (0), scalariform-opposite (1), opposite (2), opposite-alternate (3), alternate (4), none (5), with bordered pit (6), with simple pit (7) |
| C16 | Vessel distribution: solitary (0), mostly solitary (1), with some clusters and some pairs and/or radial multiples (2), mostly radial multiples and clusters with few solitary (3), none (4) |
| C17 | Vessel element in shoot: absent (0), present (1) |
| C18 | Bordered pit: absent (0), present (1) |
| C19 | Girdling leaf trace: absent (0), present (1) |
| C20 | Types of rays: homogeneous (0), heterogeneous (1), paedomorphic (2), none (3) |
| C21 | Leaf occurrence: absent (0), present (1) |
| C22 | Leaf type: microphyll (0), megaphyll (1), none (2) |
| C23 | Heterophylly: absent (0), present (1) |
| C24 | Arrangement of lateral appendages: alternate opposite (0), whorled helical (1), decussate (2), none (3), 4-rank (4) |
| C25 | Leaf organization: simple (0), compound (1), none (2) |
| C26 | Ensiform lamina: absent (0), present (1) |
| C27 | Stipule: absent (0), present (1) |
| C28 | Venation: pinnate (0), actinodromous pinnate but with basal concentration of laterals parallel (1), dichotomous other palmate (2), one-veined (3), none (4) |
| C29 | Secondary veins: brochidodromous (0), craspedodromous (1), semicraspedocromous (2), camptodromous (3), none (4) |
| C30 | Leaf margin: smooth (0), serrate (1), none (2), dissected lobate (3), |
| C31 | Leaf/pinna shape: lobate (0), lingular (1), wedge-shaped (2), linear (3), dissected lanceolate (4), fan-like (5), strap-like (6), scale-like (7), oboval (8), none (9), needle-like(A), bifid (B), orbicular (C), cordate-based (D), peltate (E), tubular (F), teret (G) |
| C32 | Midrib: absent (0), present (1) |
| C33 | Vein reticulation: absent (0), present (1) |
| C34 | Stomata occurrence: hypostomatic (0), hyperstomatic (1), amphistomatic (2), on branch (3), none (4), on sporangia (5) |
| C35 | Stomata type: anomocytic (0), paracytic (1), tetracytic (2), dicyclic (3), laterocytic (4), encyclocytic (5), desmocytic (6), diacytic (7), pericytic (8), polocytic (9), hexacytic (A), anisocytic (B), parallelocytic (C), heliocytic (D), axillocytic (E), none (F) |
| C36 | Heterospory: absent (0), present (1) |
| C37 | Gender: bisexual (0), monoecious (1), dioecious (2), none (3), androdioecious (4), polygamomonoecious (5), polygamodioecious (6), gynomonoecious (7), gynodioecious (8), andromonoecious (9) |
| C38 | Siphonogamy:absent (0), present (1) |
| C39 | Reproductive and vegetaive organ : intermixing (0), separated (1), none (2), |
| C40 | Mega- and microsporangia arrangement: proximal (0), segregated (1), none (2), |
| C41 | Reproductive parts concentrated to the distal: absent (0), present (1) |
| C42 | Reproductive organ (flower/cone) base: sessile (0), pedicellate (1), none (2) |
| C43 | Reproductive organ peripheral foliar appendage merism: numerous and helical (0), pentamerous (1), tetramerous (2), few and variable (3), none (4), trimerous (5), hexamerous (6) |
| C44 | Reproductive organ 1st cycle peripheral foliar appendage connation: absent (0), present (1), none (2) |
| C45 | Reproductive organ 2nd cycle peripheral foliar appendage connation: absent (0), present (1), none (2) |
| C46 | Petal clawed: absent (0), present (1), none (2) |
| C47 | Hypanthium:absent (0), present (1) |
| C48 | Nectary : absent (0), abaxial (1), gynoecial (2), surface (3), disc peripheral (4), foliar (5), appendage other position and roecial (6), extrafloral (7), perigonal (8), stylar (9), base stigmatic (A) |
| C49 | Microsporangia grouping: synangiate (0), distinct (1) |
| C50 | microsporangium arrangement: connate (0), free (1), none (2) |
| C51 | Heterostyly: absent (0), present (1), none (2) |
| C52 | Anther orientation: introrse (0), latrorse (1), extrorse (2), none (3) |
| C53 | Anther type: tetrasporangiate (0), bisporangiate (1), none (2), trisporangiate (3), unisporangiate (4), multisporgniate (5) |
| C54 | Microsporangium insertion: basifixed (0), dorsifixed (1), ventrifixed (2) |
| C55 | Microsporangia position relative to support structure: absent adaxial (0), abaxial (1), marginal (2), helical (3), terminal (4), peripheral (5), between bracts (6) |
| C56 | Microsporangia opening: longitudinal slit (0), valve pore (1), none (2), flap oblique slit (3), horizontal slit (4), apical slit (5), helical slit (6), decay radial slit (7) |
| C57 | Saccate pollen: bisaccate (0), monosaccate (1), none (2), pseudosaccate (3), tetrasaccate (4), trisaccate (5), |
| C58 | Microspore grouping: monad (0), dyad (1), tetrad (2), polyad (3) |
| C59 | Microspore size: > 20 um (0), < 20 um (1) |
| C60 | Tectum: absent (0), present (1) |
| C61 | Sculpture: reticulate (0), striate (1), clavate (2), rugulate (3), spinulose/scabrate (4), vermiform psilate (5), verrucate (6), foveolate (7), none (8), echinate (9), granular fossulate (A), rugose baculate (B), gemmate (C) |
| C62 | Tectal perforations: semitectate (0), tectate (1), perforate (2), imperforate (3), none (4), atectate (5), |
| C63 | Columellae :absent (0), present (1) |
| C64 | Number of ovule-enclosing unit (carpel/cupule) per reproductive organ: >5 (0), 5 (1), 4 (2), 2 (3), 1 (4), 1-5 (5), none (6), |
| C65 | Gynoecium positioning: superiort (0), only extreme baset (1), inferior basalt (2), half inferiort (3), mostly to completely inferiort (4), nonet (5) |
| C66 | Ovule-enclosing unit (carpel) stipitation: absent (0), present (1) |
| C67 | Ovule-enclosing unit (carpel) connation: distinct base of ovaries (0), connate ovaries (1), completely connate and base of styles (2), may be connate ovaries and styles completely connate (3), none (4), |
| C68 | Ovule-enclosing unit distal projection (style) position: eccentric (0), peak of ovary (1), intercalated to each other (2), absent terminal (3), decurrent (4), gynobasic (5) |
| C69 | Non-nucellar pollen receiving part (stigma) position: decurrent (0), ventral (1), apical (2), branched (3), none (4) |
| C70 | Arrangement of megasporangium complex (placentation): marginal axile-apical (0), axile-basal parietal (1),free (2), central (3), none (4), terminal peripheral adaxial (5), abaxial basal apical laminar (6), |
| C71 | Pendulous ovules: absent (0), present (1) |
| C72 | Number of megasporangium complexes per unit (carpel/cupule): many (>10) (0), few and variable (1-10) (1), one two (2), one per gynoecium (with more than one carpel/cupule) (3), none (4) |
| C73 | Number of megasporangia surrounding layers (integument): two (0), one (1), none (2) |
| C74 | Megasporangium complex orientation: orthotropous (0), anatropous (1), campylotropous (2), none amphitropous inverted (3), |
| C75 | Fruit type: follicle or septicidal capsule (0), loculicidal capsule (1), nut or samaroid nutlet (achene) (2), fleshy (berry) (3), schizocarp (4), none (5), achene (6), samara (7), drupe (8), aggregate (9), silique (A), utricle (B), caryopsis (C), pome (D) |
| C76 | Spermatozoid: absent (0), present (1) |
| C77 | Number of functional megasporangia per megasporangium: one (0), more none (1) |
| C78 | Megaspore tetrad: linear (0), tetrahedral (1), none (2), |
| C79 | Megasporangium complex enclosed: absent (0), present (1) |
| C80 | Megasporangia retained in on mother plant: absent (0), present (1) |
| C81 | Megasporangium position relative to support structure: terminal abaxial (0), adaxial marginal lateral and isolated peripheral (1), none (2) |
| C82 | Ovule-wrapping unit (cupule): absent (0), present (1) |
| C83 | Ovule-wrapping unit (cupule) opening: distal adaxial (0), proximal abaxial (1), proximal none adaxial (2), |
| C84 | Micropylar tube: absent (0), present (1) |
| C85 | Pollination through pollen tube:absent (0), present (1) |
| C86 | Archegonia in megaspore: absent (0), present (1), none (2) |
| C87 | Double fertilization: absent (0), present (1) |
| C88 | Endosperm: absent (0), present (1) |
| C89 | Angiocarpy: absent (0), present (1) |
| C90 | Vivipary: absent (0), present (1) |
| C91 | Number of cotyledon: many (0), 4-3 (1), 1 (2), none (3) |

**Table S3.** Taxa studied and GenBank accessions. A dash (-) indicates missing data, and the indicated sequences are from GenBank.

|  | *matK* | *rbcL* | ITS | *trnL* | *psbA-trnH* |
| --- | --- | --- | --- | --- | --- |
| Actinidiaceae | KP737312 | KP737327 | KP737354 | JN102166 | KP737342 |
| Adoxaceae | EF490235 | KM360613 | U88194 | AF366927 | FJ395478 |
| Aextoxicaceae | DQ182342 | HE651123 | AY590831 | AY145362 | - |
| Aizoaceae | JQ844150 | AJ235778 | AJ582944 | KF132800 | KF132892 |
| Alangiaceae | FJ644649 | DQ340449 | KP092580 | JN102141 | JF321228 |
| Alismataceae | AB040179 | HQ901563 | DQ339086 | GQ244545 | KC584951 |
| Alseuosmiaceae | AJ429378 | X87377 | EU331124 | AJ430965 | - |
| Altingiaceae | AF015650 | JX944788 | GU576669 | DQ352220 | KP095965 |
| Amaryllidaceae | JX464550 | JX903147 | AY280349 | JX464362 | KC704237 |
| Amborellaceae | DQ185522 | L12628 | - | AY145324 | - |
| Aquifoliaceae | AF542607 | KX897006 | AH007153 | FR849986 | KU198261 |
| Araceae | AY034184 | KM360996 | KJ400913 | AM933296 | KC584957 |
| Araliaceae | AB087999 | U50250 | KF727964 | AF366930 | KX347019 |
| Araucariaceae | AM920149 | U96471 | - | - | FJ17352 |
| Basellaceae | JQ844148 | MF135428 | KY968898 | - | MF143716 |
| Berberidaceae | JQ172882 | KR075627 | KC575606 | FJ626558 | JQ172934 |
| Betulaceae | AY372021 | KC482102 | - | DQ860529 | KP095389 |
| Brassicaceae | AB354274 | AY167979 | GQ202247 | AF451575 | GQ184349 |
| Brunelliaceae | EF135512 | FJ707536 | AY935891 | AF299181 | KJ426624 |
| Bruniaceae | AY490949 | L14391 | AY494012 | GQ984070 | - |
| Buxaceae | LN877447 | MF349741 | AF245412 | LN877711 | AB331298 |
| Cactaceae | HM041657 | KT164772 | AY181566 | HM041236 | KT222807 |
| Campanulaceae | EU713264 | AJ419698 | KC013706 | GQ984060 | - |
| Caprifoliaceae | GQ997392 | GQ997430 | JQ780992 | HM228560 | GU135313 |
| Caryophyllaceae | KU722867 | M77699 | KU722881 | JN589634 | GU070803 |
| Celastraceae | EU002170 | EU002277 | EU002149 | HQ393763 | GQ998178 |
| Cephalotaxaceae | AF457108 | AF456387 | JF976114 | EF660619 | EF660701 |
| Cercidiphyllaceae | AB490203 | L11673 | AF147756 | AB445242 | AB445234 |
| Chloranthaceae | AF543733 | AF332097 | D29787 | AY145334 | KP159335 |
| Clethraceae | MF350258 | L12609 | KU350161 | AJ430871 | MF348393 |
| Combretaceae | GU135057 | FJ381811 | KT279737 | LC054306 | KP319003 |
| Connaraceae | EU002174 | L29493 | AY935892 | KC428477 | KX248193 |
| Cornaceae | EU002175 | EU002276 | AY152522 | AY254238 | JF321206 |
| Crassulaceae | AF115647 | JX848453 | AB088618 | KX510105 | KP159333 |
| Crossosomataceae | DQ443456 | DQ443456 | DQ307116 | DQ307148 | KX999391 |
| Cucurbitaceae | DQ536662 | L21937 | KC922448 | HM597017 | JQ007373 |
| Cunoniaceae | JX517913 | JX572494 | AF521290 | AF299156 | - |
| Cupressaceae | AB023999 | L25756 | AY916904 | JF514857 | JF514865 |
| Cycadaceae | AF143440 | AB917042 | KC508028 | HG316537 | KP117157 |
| Cyclanthaceae | KT205206 | KT205272 | JN016949 | KJ681523 | KT205225 |
| Cyperaceae | KX036931 | AM999811 | AF168838 | AJ295759 | KR735884 |
| Davidsoniaceae | AY935930 | AF206759 | - | KC428488 | KM895252 |
| Dilleniaceae | GQ997143 | FJ860350 | AY096030 | KF953925 | KT452064 |
| *Encephalartos* | GQ203845 | AF394361 | KX130213 | - | KX152014 |
| *Ephedra* | KP788852 | FJ958029 | FJ958007 | KT033390 | - |
| Ericaceae | U61316 | L12625 | AF382730 | AF271699 | MF348522 |
| Erythropalaceae | DQ790197 | DQ790161 | DQ790230 | EF464487 | - |
| Escalloniaceae | AJ429365 | AJ419694 | U42544 | AJ430952 | KC355696 |
| Eucommiaceae | AF345323 | L01917 | HQ384682 | - | KX346918 |
| Eucryphiaceae | EU002176 | L01918 | - | KF594405 | KM895039 |
| Euphorbiaceae | LK021494 | AY794915 | AY918198 | AY794734 | GU135374 |
| Eupomatiaceae | DQ401341 | DQ861790 | AF469771 | DQ861842 | KU564673 |
| Eupteleaceae | DQ401348 | KR232440 | KR011731 | LN610908 | KR064716 |
| Fabaceae | AF203597 | KM025243 | AF156675 | AY651848 | GU396719 |
| Fagaceae | FJ185050 | KF418893 | - | AF344182 | - |
| Fouquieriaceae | EU628508 | AY725861 | AH007888 | AJ430876 | JF321292 |
| Garryaceae | KT738340 | KT740821 | JN234737 | JN234724 | KT739195 |
| Geraniaceae | KJ916373 | KJ916495 | DQ525074 | AJ884716 | KP963310 |
| *Ginkgo* | AF279806 | JQ512538 | JQ279501 | AY145323 | AF223226 |
| *Gnetum* | AY449621 | U72819 | AY449558 | AY296493 | AY849369 |
| Gomortegaceae | - | AF206773 | AF289846 | - | AF129053 |
| Gunneraceae | EU002179 | EU002279 | AF517101 | AM397168 | GQ998323 |
| Gyrostemonaceae | FJ212199 | L22440 | AF070971 | FJ212281 | - |
| Hamaelidaceae | AF013046 | KC737393 | AF094551 | DQ352357 | KC737342 |
| Hamamelidaceae | AF248617 | AY263940 | AF015654 | GU576826 | GU576755 |
| Hydatellaceae | JQ284136 | JQ284248 | JQ284173 | - | - |
| Hydrangeaceae | GU217324 | JF308660 | GU983035 | JN226776 | GU217324 |
| Hydrocharitaceae | AB088781 | AB088810 | JN578091 | - | - |
| Illiciaceae | DQ185524 | DQ182334 | AF163734 | AY145325 | - |
| Iridaceae | KC118929 | KC704850 | DQ277636 | EU939496 | KC704317 |
| Iteaceae | EF456732 | L11188 | AY231368 | - | AY138110 |
| Juglandaceae | HE966942 | KF418920 | HM049898 | AY231167 | HE966671 |
| Juncaceae | AY973527 | L12681 | KR082780 | AY344156 | HQ181837 |
| Lauraceae | KU564571 | KU564795 | HG315591 | KX674308 | KU564671 |
| Linaceae | JX661951 | FJ169596 | KX147539 | FJ160887 | GQ845215 |
| Loasaceae | KY286710 | U17876 | KY286627 | KY286979 | JF321277 |
| Loganiaceae | KJ815653 | Z68826 | DQ358879 | AJ430910 | - |
| Lythraceae | GU228449 | L10223 | - | AY905482 | JQ730675 |
| Magnoliaceae | AY00900 | AY008933 | EU593549 | AY305819 | HM236892 |
| Melastomataceae | KP093302 | GQ436728 | KY798014 | GQ265885 | JF708230 |
| Monimiaceae | AJ247183 | AF040664 | GU177689 | AF129041 | AF129068 |
| Myricaceae | KF419021 | KF418924 | KP092762 | DQ501450 | KP095418 |
| Myrothamnaceae | AM396507 | AF060707 | AF094555 | AM397169 | EU213839 |
| Myrtaceae | AF368206 | AB537496 | - | HQ287716 | KP142183 |
| Najadaceae | HM240463 | U03731 | KT596618 | KT596674 | AB331276 |
| Nepenthaceae | AF315891 | L01936 | AB675865 | JX042575 | MF348377 |
| Neuradaceae | - | U06814 | KJ004316 | - | - |
| Nitrariaceae | - | DQ267158 | DQ267176 | DQ267166 | KP087763 |
| Nymphaeaceae | DQ185527 | LC219121 | JF805747 | AF543729 | KC525065 |
| Nyssaceae | JF308675 | JF308651 | EU734443 | EU734496 | JF321237 |
| Ochnaceae | KF263283 | KF263405 | KF263218 | HQ599792 | - |
| Olacaceae | GQ997871 | GQ997898 | DQ333869 | DQ340620 | GQ997883 |
| Onagraceae | HM851006 | EU255778 | GU176556 | GU176588 | - |
| Orchidaceae | AF263677 | AF074211 | AY273748 | AY273651 | FJ460374 |
| Oxalidaceae | EU002186 | EU002282 | EU436875 | JN639556 | GQ998541 |
| Papaveraceae | KM364817 | U86629 | - | AY328253 | - |
| Parnassiaceae | JF954971 | JN105081 | JF811045 | DQ860575 | JF802315 |
| Penaeaceae | AY151582 | AY151706 | AM235864 | - | AY151675 |
| Penthoraceae | KC988293 | AF190431 | KC988286 | X71987 | DQ006173 |
| Philydraceae | EU499296 | AY298834 | - | AJ387741 | - |
| Phytolaccaceae | AY042631 | FJ860398 | KU377334 | KM261955 | MF143636 |
| Picramniaceae | - | AF127025 | - | - | KJ426870 |
| Pinaceae | EF440492 | EF440570 | - | EF440530 | FJ493294 |
| Piperaceae | DQ882215 | EF591363 | KF924100 | EU519779 | KM055199 |
| Pittosporaceae | DQ133794 | HM850262 | HM116994 | AJ430960 | KM895297 |
| Platanaceae | EU642711 | AF081073 | AY706029 | AY145358 | KP402512 |
| Poaceae | KM487287 | AY836162 | - | JX456355 | HQ894425 |
| Podocarpaceae | AF228112 | AF249648 | AB023989 | EF660630 | JN041002 |
| Polemoniaceae | L34205 | KX397893 | AF167227 | EF433257 | KJ686557 |
| Portulacaceae | EU834751 | M62568 | JF508549 | - | MF143649 |
| Primulaceae | HQ535964 | HQ619773 | - | KM198463 | - |
| Proteaceae | AY823204 | JX571862 | - | AF482140 | - |
| Ranunculaceae | EU827661 | DQ099444 | FJ424225 | KP900425 | KY235737 |
| Rhamnaceae | GQ434248 | KP121449 | DQ146573 | KR083150 | HG765028 |
| Rhizophoraceae | JX661962 | JX664070 | AF130332 | AY947404 | JX663300 |
| Rhoipteleaceae | U92852 | AF017687 | AF303800 | AY147081 | KF201474 |
| Rosaceae | DQ860466 | JQ391365 | JQ392451 | DQ863238 | JQ390723 |
| Rubiaceae | KJ815700 | FJ493349 | - | KJ815420 | MF348541 |
| Sabiaceae | HM755905 | HM755932 | KM092335 | KP900465 | HM755920 |
| Sapindaceae | KP675825 | FN599461 | KP675785 | JN102157 | KJ687156 |
| Sarraceniaceae | JQ619005 | KR819588 | DQ098117 | AJ430886 | MF348369 |
| Saxifragaceae | KC737243 | KC737395 | KC004032 | JN102287 | JF708213 |
| Scrophulariaceae | KR424749 | EF544598 | EF363673 | DQ856494 | KR424763 |
| Solanaceae | EF537313 | AF397102 | KP006659 | EU603443 | EF537224 |
| Staphyleaceae | EU002189 | EU002285 | KR349376 | KX999498 | KX999401 |
| Symplocaceae | AF380108 | AY725865 | MF171080 | AF534682 | EU769795 |
| Taccaceae | AY973837 | JF944589 | JF978861 | JN850578 | JN047324 |
| Taxodiaceae | AB030115 | JQ512531 | AB023983 | AY727216 | JQ512286 |
| Tetracarpaeaceae | L34154 | L11207 | EF178790 | - | - |
| Theaceae | - | AF380058 | MF171105 | HM100596 | HM100528 |
| Thymelaeaceae | KU244186 | GQ436619 | KX024773 | GU736358 | KU244056 |
| Trapaceae | KX526694 | L10226 | FM887019 | AY905491 | - |
| Trochodendraceae | AF543751 | GQ998840 | AF192799 | AY145360 | GQ998829 |
| Tropaeolaceae | FM179931 | L14706 | JN115053 | AB043665 | - |
| Turneraceae | JX661965 | DQ123398 | DQ521285 | JQ723403 | JX663303 |
| Ulmaceae | KC539623 | KC539693 | KC539589 | KC539738 | - |
| Velloziaceae | JX286710 | AY149358 | JN016957 | AF293095 | JN017061 |
| Vochysiaceae | GQ982128 | AM235665 | DQ787415 | - | KX249560 |
| *Welwitschia* | AF280996 | AF394335 | U50740 | AY513733 | AY849370 |
| Winteraceae | EU669474 | EU669518 | AY526318 | EU669556 | FJ539202 |
| *Zamia* | GQ203859 | JQ770263 | AJ287364 | - | JQ770199 |

**Figure S1.**
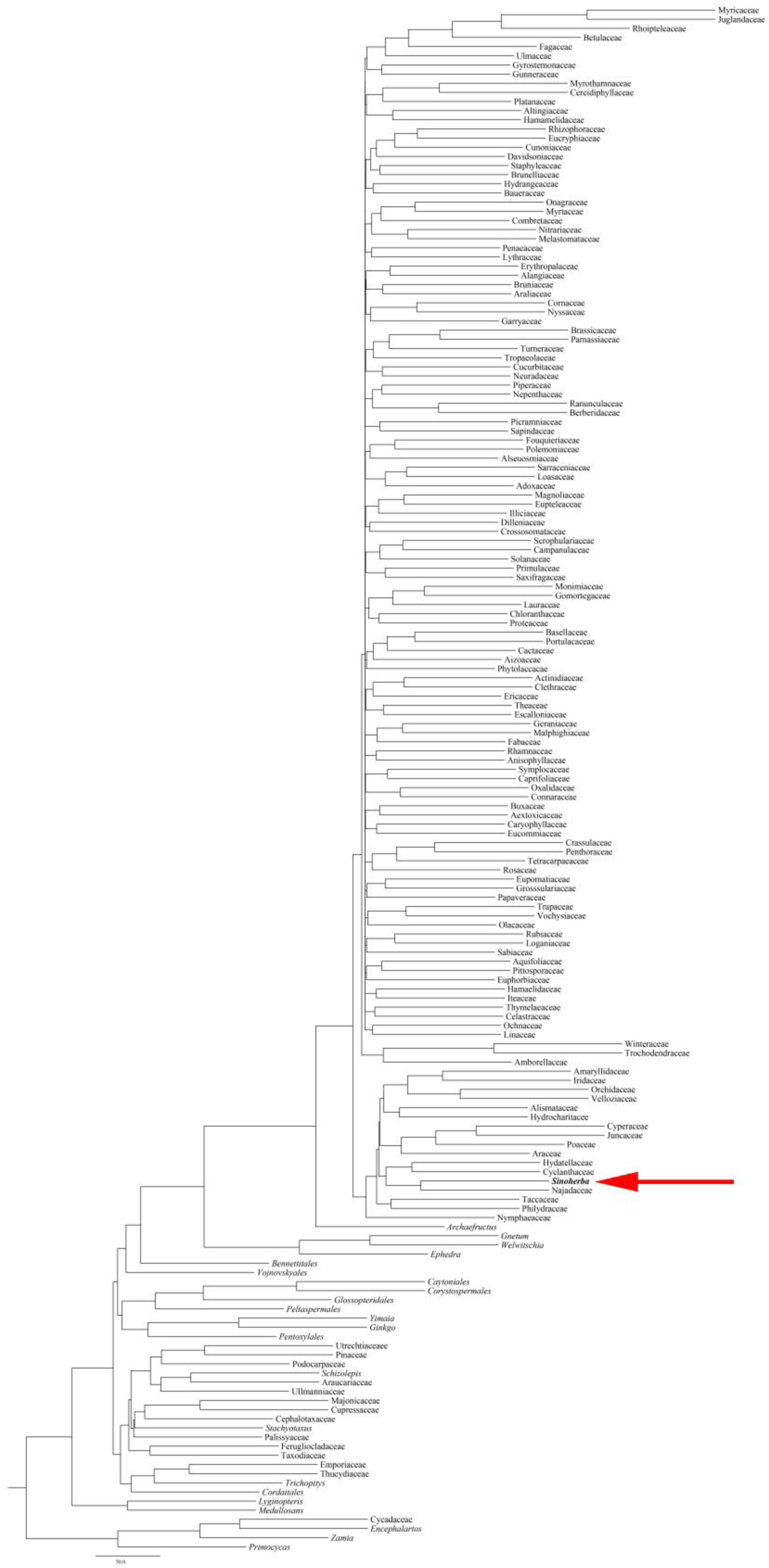
Phylogenetic relationships of the morphological characteristic score information. The numbers near the nodes are bootstrap percentages. “∗” indicates that the node is 100% supported. “←” indicates the phylogenetic position of *Sinoherba.*

**Figure S2.**
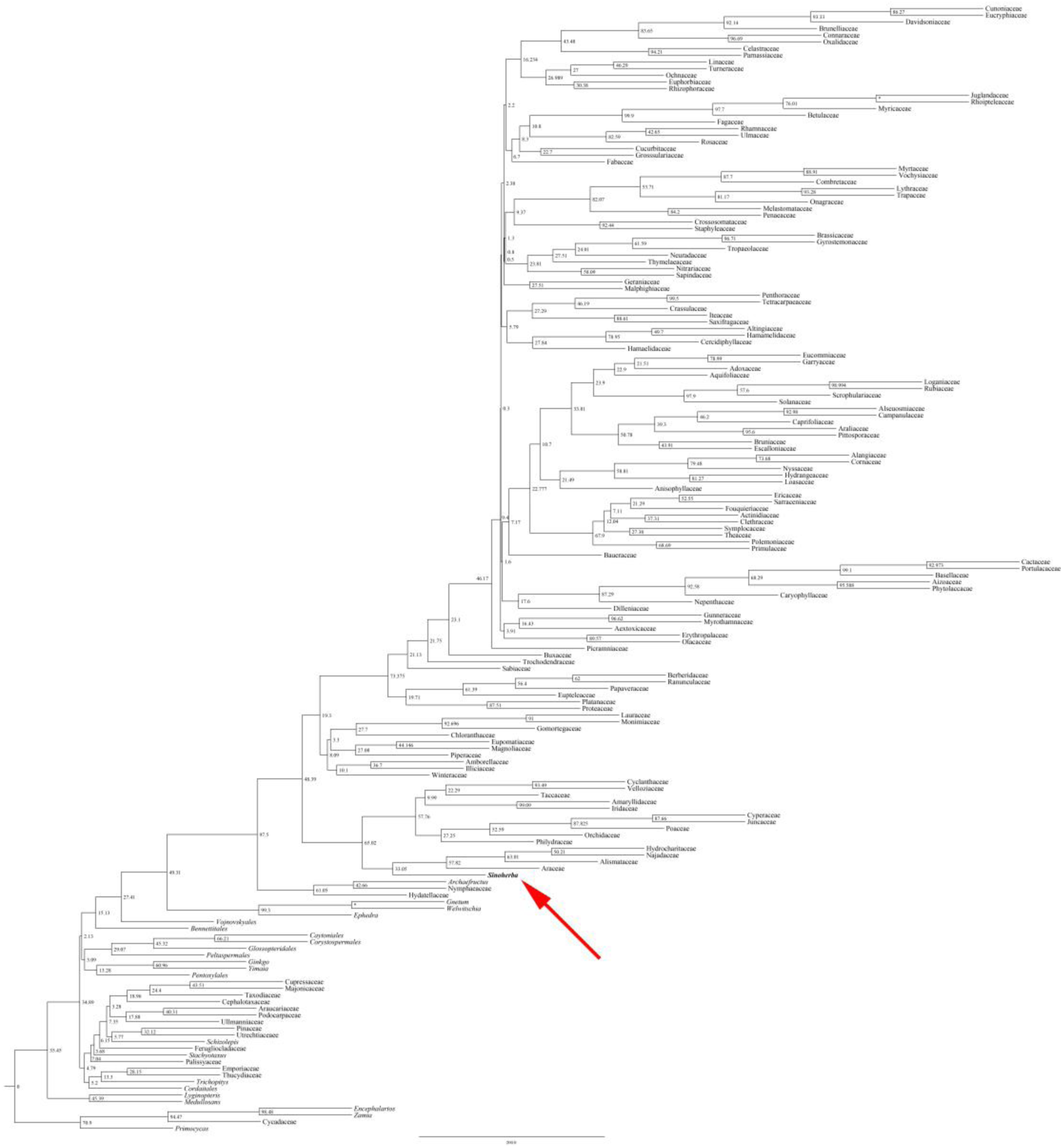
Phylogenetic relationships based on DNA sequence (*rbcL*, *matK*, *trnL*-*F*, *psbA*-*trnH* and ITS) and morphological characteristic score information of *Sinoherba.* The numbers near the nodes are bootstrap percentages. “∗” indicates that the node is 100% supported. “←” indicates the phylogenetic position of *Sinoherba.*

